# Molecular characterization of differences between the tomato immune receptors Fls3 and Fls2

**DOI:** 10.1101/2020.02.04.934133

**Authors:** Robyn Roberts, Alexander E. Liu, Lingwei Wan, Annie M. Geiger, Sarah R. Hind, Hernan G. Rosli, Gregory B. Martin

## Abstract

Plants mount defense responses by recognizing indications of pathogen invasion, including microbe-associated molecular patterns (MAMPs). Flagellin from the bacterial pathogen *Pseudomonas syringae* pv. tomato (*Pst*) contains two MAMPs, flg22 and flgII-28, that are recognized by tomato receptors Flagellin sensing 2 (Fls2) and Flagellin sensing 3 (Fls3), respectively. It is unknown to what degree each receptor contributes to immunity and if they promote immune responses using the same molecular mechanisms. Characterization of CRISPR/Cas9-generated *Fls2* and *Fls3* tomato mutants revealed that the two receptors contribute equally to disease resistance both on the leaf surface and in the apoplast. However, striking differences were observed in certain host responses mediated by the two receptors. Compared to Fls2, Fls3 mediated a more sustained production of reactive oxygen species (ROS) and an increase in transcript abundance of 44 tomato genes, with two genes serving as reporters for Fls3. Fls3 had greater *in vitro* kinase activity and interacted differently with the *Pst* effector AvrPtoB as compared to Fls2. Using chimeric Fls2/Fls3 proteins, we found that no receptor domain was solely responsible for the Fls3 sustained ROS, suggesting involvement of multiple structural features. This work reveals differences in the immunity outputs between Fls2 and Fls3, suggesting they use distinct molecular mechanisms to activate pattern-triggered immunity in response to flagellin-derived MAMPs.

## Introduction

*Pseudomonas syringae* pv. tomato (*Pst*) causes bacterial speck disease on tomato, resulting in small, necrotic lesions on leaves, stems, fruit, and flowers (Jones, 1991). The tomato-*Pst* pathosystem serves as a model for studying bacterial pathogenesis and plant immunity (Oh and Martin, 2011; Pedley and Martin, 2003). The virulence of *Pst* is primarily determined by a suite of 36 type III effectors which are delivered into the plant cell during the infection process. Two effectors, AvrPto and AvrPtoB, play major roles in interfering with plant immune responses and promote enhanced multiplication of *Pst* in the leaf apoplast (Buell et al., 2003; Cunnac et al., 2011; Kvitko et al., 2009; Lin and Martin, 2005; Martin, 2012).

Plants detect pathogens by recognizing conserved microbe-associated molecular patterns (MAMPs) via pattern recognition receptor (PRR)-triggered immunity (PTI), or by recognizing pathogen effectors through nucleotide-binding oligomerization domain like receptors (NLR)-triggered immunity (NTI). For *Pst*, the motility-associated protein flagellin contains two MAMPs, flg22 and flgII-28, that are recognized by the PRRs Flagellin sensing 2 (Fls2) and Flagellin sensing 3 (Fls3), respectively (Gomez-Gomez and Boller, 2000; Hind et al., 2016; Robatzek et al., 2007). Upon recognition of these and other MAMPs, a suite of molecular events occurs to promote defense, including the production of reactive oxygen species (ROS), activation of a mitogen-activated protein kinase (MAPK) cascade, transcriptional reprogramming, callose deposition at the cell wall, stomatal closure, and calcium fluxes (Couto and Zipfel, 2016; Li et al., 2016). While PTI has generally been associated with a moderate inhibition of pathogen growth (∼10-fold), a recent study shows that flagellin-mediated PTI plays a major role in immunity on the leaf surface for some tomato accessions (decreasing bacterial populations by ∼150-fold) (Roberts et al., 2019b).

Fls2 and Fls3 bind flg22 and flgII-28 through their extracellular leucine-rich repeat (LRR) domain. In Arabidopsis, upon binding flg22, FLS2 associates with the co-receptor BRI1-ASSOCIATED RECEPTOR KINASE (BAK1) and both FLS2 and BAK1 are transphosphorylated to initiate downstream signaling (Couto and Zipfel, 2016; Saijo et al., 2018; Sun et al., 2013). Tomato has two *Fls2* genes, *Fls2.1* (Solyc02g070890) and *Fls2.2* (a paralog of *Fls2.1* located 3.8 kB away from *Fls2.1*) (Solyc02g070910). *Fls2.1* appears to encode the only functional Fls2 in tomato, as a mutation in *Fls2.1* causes a complete loss of flg22 recognition (Jacobs et al., 2017). Previous work has shown that not all solanaceous species have Fls3. While tomato, potato, and pepper respond to flgII-28, *Nicotiana benthamiana* and petunia do not (Cai et al., 2011; Clarke et al., 2013; Hind et al., 2016). A previous study found some similarities between the components involved in Fls2 and Fls3 signaling in tomato, including that Fls3 can interact with Arabidopsis BAK1 upon flgII-28 induction (Hind et al., 2016). Similar to *Fls2.1, Fls3* gene expression is induced in leaves after treatment with flg22, flgII-28, or DC3000Δ*avrPto*Δ*avrPtoB* (Hind et al., 2016; Rosli et al., 2013). After flgII-28 treatment, there is an increase in the phosphorylation of MAPKs in protoplasts transfected with *Fls3* (Hind et al., 2016). *N. benthamiana* plants silenced for *Bak1* have a reduced flgII-28 ROS response compared to control plants, and Fls3 and Arabidopsis BAK1 interact upon flgII-28 treatment when Fls3 and BAK1 are co-expressed in *N. benthamiana* leaves (Hind et al., 2016). Together, these observations suggest there may be similar factors involved in Arabidopsis Fls2 and tomato Fls3 signaling, but further analysis is needed to determine the molecular mechanisms of Fls3.

Fls2 and Fls3 belong to a family of non-RD kinases due to their lack of conserved arginine (R) and aspartate (D) residues in the activation loop. In Arabidopsis, FLS2 has been shown to have weak autophosphorylation activity that requires the presence of the entire FLS2 intracellular domain, including the inner juxtamembrane domain and the kinase domain (Cao et al., 2013; Gomez-Gomez et al., 2001; Lu et al., 2010; Xiang et al., 2008). Chimeric constructs containing different domains of PRRs have been made to study which domains are important for receptor functions (Albert and Felix, 2010; Albert et al., 2010; Brutus et al., 2010; Hohmann et al., 2018; Holton et al., 2015; Kouzai et al., 2013; Mueller et al., 2012; Wu et al., 2019; Zhou et al., 2019). Chimeric constructs combining the ectodomain of Arabidopsis receptor-like kinase EFR (which detects the bacterial MAMP Ef-Tu) with the intracellular domain of cell wall-associated kinase AtWAK1 (which recognizes oligogalacturonides released from the cell wall) results in a functional chimeric protein that recognizes Ef-Tu and activates plant defenses. However, combining the AtWAK1 ectodomain with the EFR intracellular domain did not result in sufficient defenses, as it did not cause a significant reduction in bacterial growth compared to the negative control (Brutus et al., 2010). Combining the FLS2 ectodomain with the EFR intracellular domain also resulted in a functional protein (Brutus et al., 2010). A chimeric cross-species protein combining the Arabidopsis EFR extracellular domain with the transmembrane and intracellular domains of rice Xa21 (an LRR-RLK that recognizes the sulfated protein RaxX of *Xanthamonas oryzae* pv. oryzae and encodes resistance) was also functional (Holton et al., 2015). Chimeric constructs swapping Arabidopsis FLS2 and tomato Fls2.1 LRRs aided the authors in finding specific LRR repeats in tomato that are responsible for recognizing the flg15 peptide, which is not recognized by Arabidopsis Fls2 (Mueller et al., 2012; Robatzek et al., 2007).

NTI is activated upon the recognition of pathogen effectors and causes a suite of molecular events that includes the activation of a MAPK cascade, transcriptional reprogramming, and localized, controlled cell death (hypersensitive response, HR) that is typically associated with a significant inhibition of pathogen growth (∼100-1000 fold). The *Pst* effectors AvrPto and AvrPtoB are recognized by the cytoplasmic kinase Pto, which acts with the NLR Prf to activate the immune response and results in cell death (Dong et al., 2009; Gutierrez et al., 2010; Lin and Martin, 2007; Martin, 2012; Mathieu et al., 2014; Mucyn et al., 2009; Salmeron et al., 1996; Xing et al., 2007). AvrPto and AvrPtoB are unrelated effectors, but both can bind to Pto. Certain variants of AvrPtoB are also bound by the Fen kinase which is closely related to Pto (Martin, 2012; Martin et al., 1993; Mathieu et al., 2014; Rosebrock et al., 2007). AvrPtoB contains a Pto-interacting domain (PID) near the N-terminus (within residues 1-307), a Fen-interacting domain (FID) in the middle (residues 307-387), and an E3 ubiquitin ligase at the C-terminus. The E3 ligase degrades Fen when Fen is bound to the FID, but because Pto can bind to the FID or PID, Pto can evade E3 ligase degradation by binding to the PID. Deleting the E3 ligase from AvrPtoB (AvrPtoB_1-387_) allows both Pto and Fen to bind the FID, resulting in a strong cell death response (Mathieu et al., 2014). Using a MAPK suppression assay, AvrPtoB_1-359_ was determined to be the minimally biologically functional region of AvrPtoB, which eliminates the E3 ligase and truncates part of the FID but is still able to bind to Fen (Cheng et al., 2011; Martin, 2012; Martin et al., 1993; Mathieu et al., 2014).

Importantly, NTI may have evolved in response to effectors blocking the PTI pathway to aid pathogen virulence. AvrPtoB binds to the Arabidopsis BAK1 kinase domain (BAK1-KD) and inhibits BAK1 kinase activity *in vitro,* likely by competing for the BAK1 substrate binding site via an overlapping binding domain with Fen (AvrPtoB_250-359_) and does not require the E3 ligase domain (Cheng et al., 2011; Rosebrock et al., 2007; Shan et al., 2008). Mutation of AvrPtoB_250-359_ to AvrPtoB_250-359_(R271A/R275A) results in a loss of interaction of AvrPtoB_250-359_ and BAK1-KD by abolishing the interaction of R271 and R275 with the phosphate group of T450 in BAK1, which is required to maintain BAK1 in its active form (Cheng et al., 2011).

Here, we investigated some of the similarities and differences in the immunity outputs between Fls2 and Fls3 in tomato and present data suggesting there are some important differences in the molecular signaling between these two flagellin-sensing receptors in tomato.

## Results

### Fls3 and Fls2 each contribute to disease resistance in tomato

We used the CRISPR/Cas9 system to develop tomato lines that were insensitive to the peptides flgII-28, flg22, or both flgII-28 and flg22. In the background of Rio Grande-prf3 (RG-prf3, which has a mutation in *Prf* that eliminates the defense responses associated with the bacterial effectors AvrPto and AvrPtoB), we generated mutations in *Fls3* alone (ΔFls3), *Fls2.1* alone (ΔFls2.1), both *Fls2.1* and its paralog *Fls2.2* together (ΔFls2.1/2.2), and *Fls2.1, Fls2.2,* and *Fls3* together (ΔFls2.1/2.2/3) and verified the presence of mutations using Sanger sequencing (Table S1, Fig. S1a, and Supplementary Methods). The CRISPR/Cas9-generated mutations resulted in a 1 base pair insertion (+1bp) in the gRNA region of *Fls3* for ΔFls3, a -7bp deletion (-7bp) in *Fls2.1* for ΔFls2.1, a -7bp deletion *Fls2.1* and a 1bp insertion (+1bp) in *Fls2.2* for ΔFls2.1/2.2, and a +1bp insertion in each of the three *Fls2.1, Fls2.2,* and *Fls3* genes for ΔFls2.1/2.2/3. The loss of flgII-28 or flg22 peptide recognition was confirmed in each mutant line using ROS assays (Fig. S1b and Table S1).

To test the contribution of Fls3 and Fls2 to flagellin recognition on the leaf surface and mounting of defense responses, the four mutant lines and wild-type RG-prf3 were dip inoculated with *P. syringae* pv. tomato (*Pst*) DC3000 strains deleted for *avrPto* and *avrPtoB* (DC3000Δ*avrPto*Δ*avrPtoB*) or *avrPto, avrPtoB,* and *fliC* (DC3000Δ*avrPto*Δ*avrPtoB*Δ*fliC*). Bacterial growth in leaves was measured two days after inoculation. In the wild-type, ΔFls2.1, ΔFls2.1/2.2 and ΔFls3 lines we observed significant differences in bacterial growth between the two strains, suggesting that all these lines are able to recognize flagellin. However, in the ΔFls2.1/2.2/3 line there was no difference in bacterial growth between DC3000Δ*avrPto*Δ*avrPtoB* and DC3000Δ*avrPto*Δ*avrPtoB*Δ*fliC* (Fig. 1A).

**Figure 1.**
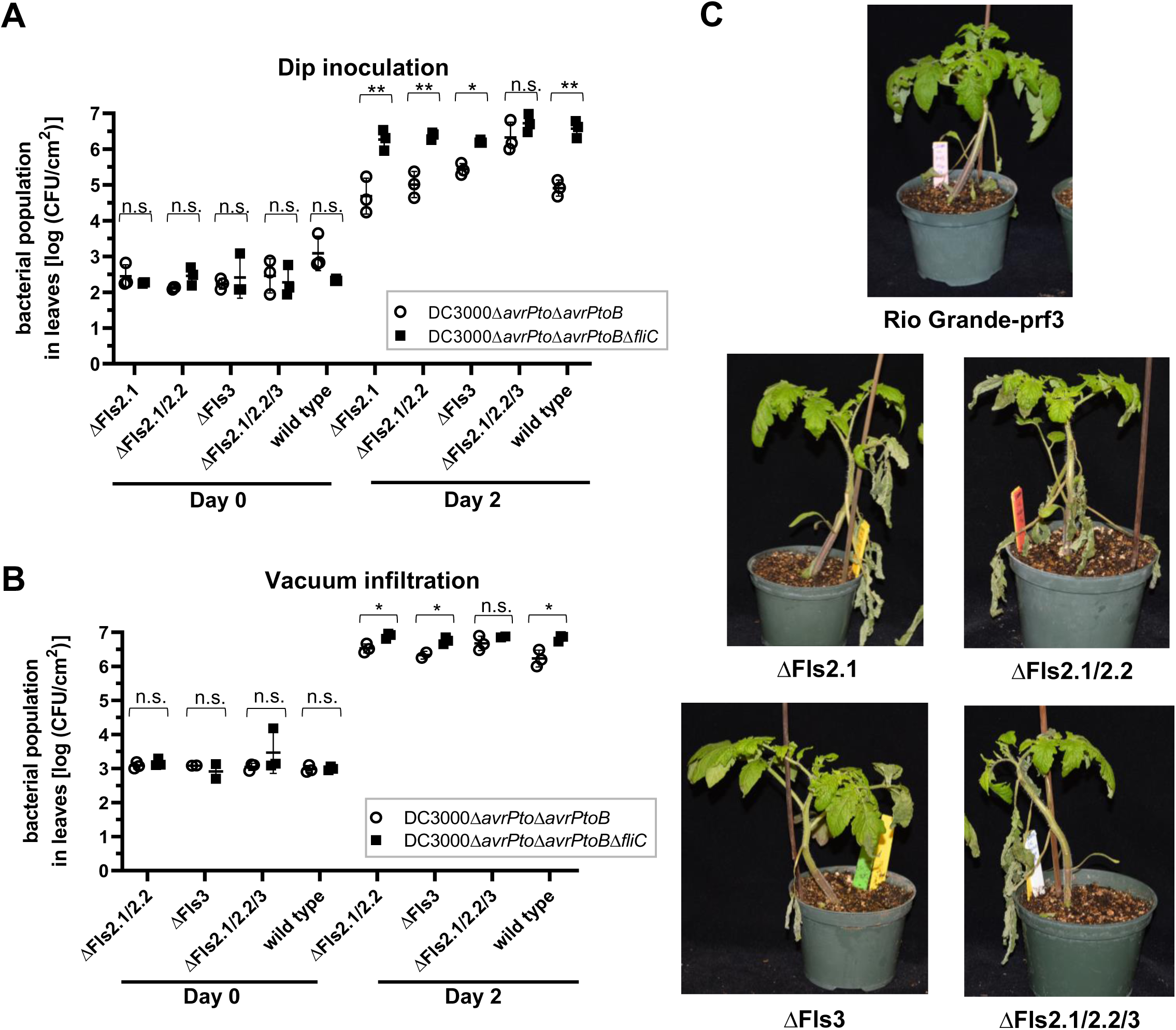
Both Fls2 and Fls3 contribute to disease resistance on the leaf surface and in the leaf apoplast in tomato. Bacterial populations in tomato leaves in CRISPR/Cas9-generated mutants of *Fls2.1* (ΔFls2.1), *Fls3* (ΔFls3), *Fls2.1* and *Fls2.2* (ΔFls2.1/2.2), or *Fls2.1, Fls2.2,* and *Fls3* (ΔFls2.1/2.2/3) or Rio Grande-prf3 (RG-prf3) were measured 0 and 2 days after **(A)** dip-inoculating plants in bacterial suspensions (1 x 10^8^ cfu/mL), or **(B)** vacuum infiltrating plants with bacterial suspensions (1 x 10^4^ cfu/mL). Shown are the means of three individual plants indicated as separate points, and horizontal lines are the means of the three plants +/- s.d.. Statistical significance was determined by pairwise t-test, where \**P*<0.05, \*\**P*<0.01, or *P*>0.05 (not significant, n.s.). **(C)** Photos of CRISPR/Cas9-generated mutant plants vacuum infiltrated with DC3000*ΔavrPtoΔavrPtoB* at 1×10^4^ CFU/mL, taken 1 week post inoculation. Experiments were repeated three times with similar results and data are representative of a single replicate. *See also Figure S1 and Tables S1–S2*.

We next tested whether Fls3 and Fls2 contribute to disease resistance in the leaf apoplast by vacuum infiltrating DC3000Δ*avrPto*Δ*avrPtoB* and DC3000Δ*avrPto*Δ*avrPtoB*Δ*fliC* into ΔFls2.1/2.2, ΔFls3, and ΔFls2.1/2.2/3 plants and the wild-type line as a control and measuring bacterial growth two days later. Similar to dip-inoculated plants, we observed differences in bacterial growth between the two strains for all of the lines except for ΔFls2.1/2.2/3. However, the differences in growth observed for the vacuum-inoculated ΔFls2.1/2.2, ΔFls3, and RG-prf3 were much smaller compared to the dip-inoculated plants, with about a 10-fold difference in bacterial growth when vacuum infiltrated versus about a 130-fold difference when dip inoculated (Fig. 1B). This suggests that Fls3 and Fls2 act both on the leaf surface and in the apoplast, but the effect is greater on the leaf surface. We also observed that, compared to RG-prf3, speck symptoms were more severe on the plants that are unable to recognize flg22 and/or flgII-28 (ΔFls2.1, ΔFls2.1/2.2, ΔFls3, and ΔFls2.1/2.2/3). Symptoms were similar on ΔFls2.1, ΔFls2.1/2.2, and ΔFls3 plants, and were slightly more severe on ΔFls2.1/2.2/3 plants (Fig. 1C). Together, these observations suggest that Fls3 and Fls2 both make important contributions to *Pst* resistance in tomato.

### The ROS response induced by flgII-28 is more sustained compared to that induced by flg22

We previously observed a more sustained ROS response mediated by Fls3 compared to Fls2 in an accession of the wild species *S. pimpinellifolium* and hypothesized this might indicate the two PRRs use different molecular mechanisms (Hind et al., 2016). To follow up on this observation, we tested whether tomato (RG-prf3) had a sustained ROS response to flgII-28 compared to flg22. As previously observed, we saw a sustained ROS response for flgII-28 compared to flg22 (Fig. 2A and S2AB). While the amplitude of the flgII-28 and flg22 responses was similar, the flgII-28 ROS response continued for up to 100 minutes versus 60 minutes for flg22 (Fig. S2A). To see the generality of this sustained response, we tested another tomato accession (CR293), a tomato heirloom (Galina), and a tomato breeding line (OH05-8144) and found that the flgII-28 response was sustained in all of these lines compared to flg22 (Fig. S2B).

**Figure 2.**
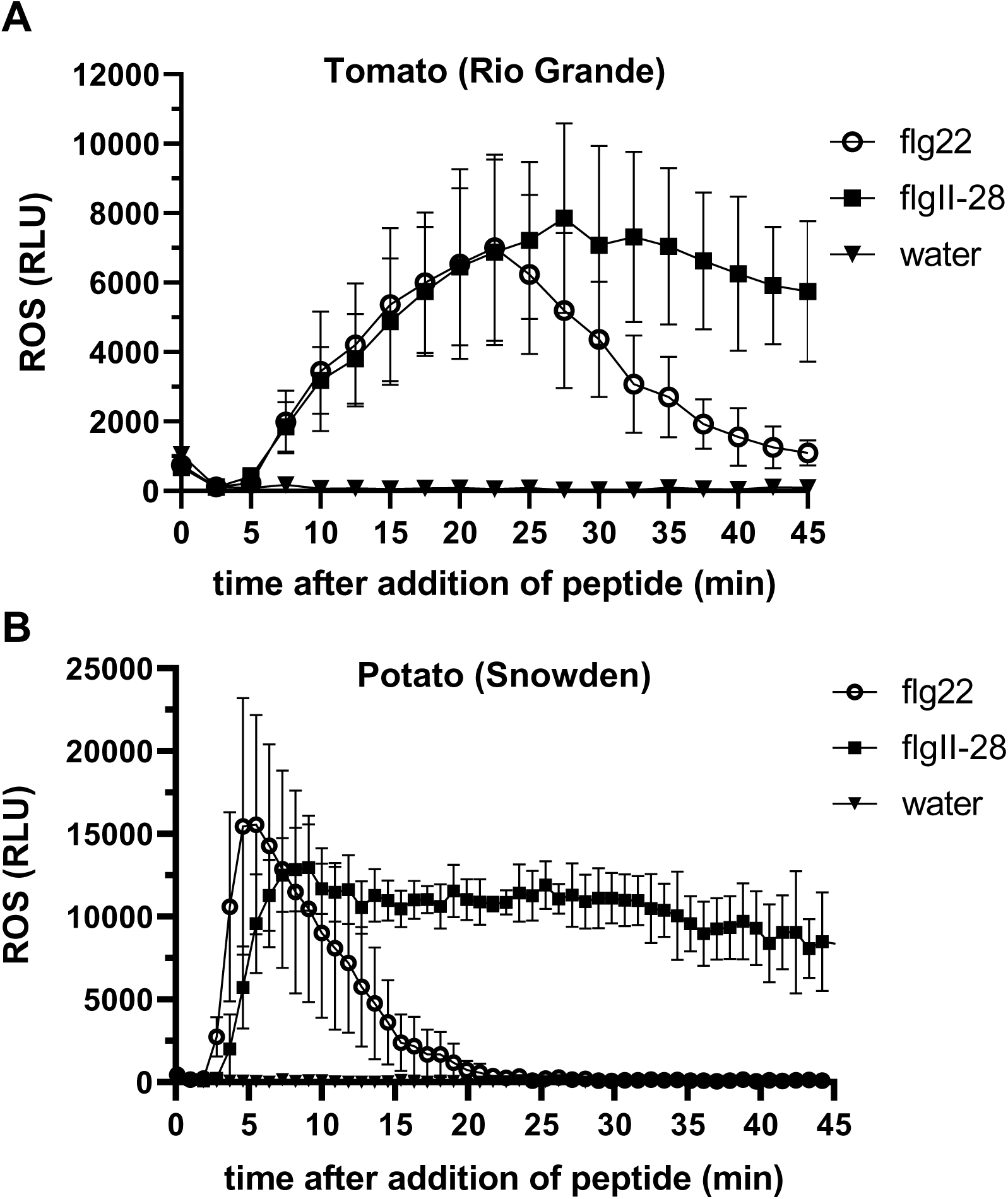
The flgII-28 ROS response is sustained in both tomato and potato. Oxidative burst produced over 45 minutes in response to 100 nM flgII-28 or flg22 peptide or water in **(A)** tomato variety Rio Grande-prf3, or **(B)** potato variety Snowden. ROS was measured in relative light units (RLU) and results shown are means +/-s.d. (n=4 for tomato, n=3 for potato) and are representative of three independent experiments. *See also Figures S2 and S3*.

To investigate if this sustained ROS response was specific to tomato, we also tested the flgII-28 ROS response in another solanaceous species, potato (*Solanum tuberosum*), for which it has been shown that flgII-28 is a major MAMP but not how sustained the associated ROS production was (Clarke et al., 2013; Moroz and Tanaka, 2019). We found that two potato accessions, Snowden and Dakota Crisp, both had a sustained ROS response for flgII-28 compared to flg22, though the amplitude of the response to both MAMPs was similar (Fig. 2B and S2C). A previous study observed a sustained ROS response for pepper *(Capsicum annuum* cv. ‘Jalapeno Early’) (Clarke et al., 2013). Because only one variety of eggplant (MM643, *Solanum melongena*) was previously tested and shown non-responsive to flgII-28 (Clarke et al., 2013), we tested the eggplant variety Shikou for its flgII-28 response and found that it did not respond to flgII-28 (Fig. S3). Together, our data and previous reports suggest that the flgII-28 extended ROS response is conserved in all solanaceous species that respond to flgII-28.

### Fls3 specifically regulates expression of a set of defense-related genes in tomato leaves

Reporter genes are useful for monitoring the activity of specific signaling pathways. From data generated in a previous RNA-Seq analysis of flagellin-induced genes in tomato (Pombo et al., 2017; Rosli et al., 2013) we identified 44 genes whose transcript abundance is significantly increased 6 hours after applying flgII-28, but not flg22 (Table S2). Gene ontology enrichment (GO term) analysis revealed that many of these genes were defense-related, including terms related to proteolysis, peptidase and hydrolase activity, and metabolism (Table S3). Among these genes, the two most highly induced encode a BHLH transcription factor (*BHLH*, Solyc03g114230) and a UDP-glucosyltransferase (*UGT*, Solyc09g098080). To test whether these genes might serve as specific reporters for Fls3 activity, we used RT-qPCR to estimate gene expression in leaves of the two most highly induced genes, *BHLH* and *UGT,* after infiltrating 1μM flgII-28 or flg22, or water as a control. We found that transcript abundance of both *BHLH* and *UGT* was significantly increased 6 hours after flgII-28 treatment compared to flg22 (Fig. 3). After flgII-28 treatment, *BHLH* gene expression was induced by 117-fold and *UGT* was induced about 303-fold relative to the water treatment. Comparatively, flg22 induced expression of *BHLH* by 3-fold and *UGT* by 12-fold. These data suggest that *BHLH* and *UGT* can serve as specific reporters for the Fls3 response to flgII-28.

**Figure 3.**
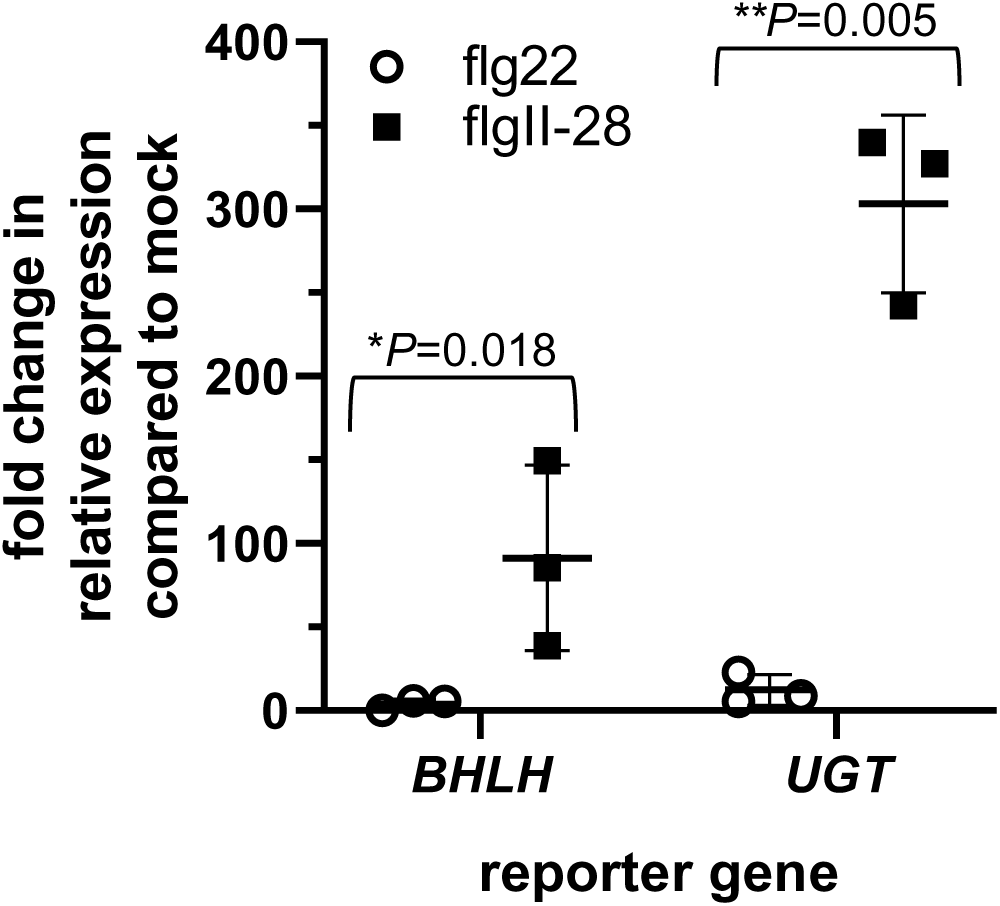
Expression of four tomato genes is induced by the Fls3 pathway but not the Fls2 pathway. qRT-PCR of the two most highly induced genes (BHLH transcription factor (BHLH, Solyc03g114230) and a UDP-glucosyltransferase (UGT, Solyc09g098080)) was performed 6 hr after infiltrating 1μM flg22 or flgII-28 peptide or water, and normalized to an internal control (ARD2). Data shown are the average fold change in expression compared to the water control of three individual plants from a single experimental replicate shown as separate points, and horizontal lines are the means of the three plants +/- s.d.. Significance was determined by pairwise t-test and *P* values are shown above each pairwise comparison. Asterisks represent significance of \**P*<0.05 or \*\**P<*0.01. Results are representative of three independent experiments.

### Fls3 and Fls2 are inhibited differently by the bacterial effector AvrPtoB

The effector AvrPtoB blocks PTI by interacting with Bak1 and inhibits Bak1 kinase activity through a mechanism involving the AvrPtoB_250-359_ region (Cheng et al., 2011). We have observed that the Fls3 ROS output is similar between Fls3 natively expressed in tomato and tomato Fls3 that is transiently overexpressed in *N. benthamiana* leaves, making transient expression a useful system to study Fls3 (this study and (Hind et al., 2016)). In Arabidopsis, AvrPtoB_1-359_ binds to the co-receptor BAK1 to inhibit PTI and promote bacterial growth. The mutant, AvrPtoB_1-359_(R271A/R275A), is unable to bind to BAK1, and therefore PTI is not suppressed in this pathway. Therefore, if Fls3 and Fls2 both interact with the BAK1 ortholog in tomato (SERK3A/SERK3B), we expect that ROS would be suppressed when Fls3 or Fls2 is co-expressed with AvrPtoB_1-359_, but not with AvrPtoB_1-359_(R271A/R275A). We tested this by transiently co-expressing Fls3 and Fls2 with AvrPtoB_1-359_, AvrPtoB_1-359_(R271A/R275A), or YFP as a control in *N. benthamiana* that was silenced for *Prf* using a hairpin construct (hpPrf) (to avoid detection of AvrPtoB by Pto/Prf) and measured ROS after the addition of 100 nM flgII-28 or flg15 peptide, 60 hours after infiltration of the various constructs. We used the flg15 peptide instead of flg22 to circumvent detection of endogenous *N. benthamiana* Fls2 (flg15 is recognized by tomato Fls2 and not by *N. benthamiana* (Robatzek et al., 2007)). We observed that Fls3 ROS production in response to flgII-28 peptide was similarly inhibited by AvrPtoB_1-359_ as the Fls2 ROS response to flg15 (Fig. 4A). We then tested whether co-expressing Fls3 and Fls2 with the mutant AvrPtoB_1-359_(R271A/R275A) would fail to suppress PTI in both RLKs, which would lead to a recovery of the Fls3 or Fls2 ROS response. As expected, when Fls2 was co-expressed with AvrPtoB_1-359_(R271A/R275A) ROS was not inhibited, and we observed similar ROS production as co-infiltrating with the YFP control. Unexpectedly, co-expression of Fls3 with AvrPtoB_1-359_(R271A/R275A) still inhibited Fls3 ROS. Protein expression was confirmed via immunoblot for all proteins except for AvrPtoB_1-359_(R271A/R275A) (Fig. S4A); however this protein is likely expressed at low levels since it was able to suppress Fls3-associated ROS production in Fig. 4A.

**Figure 4.**
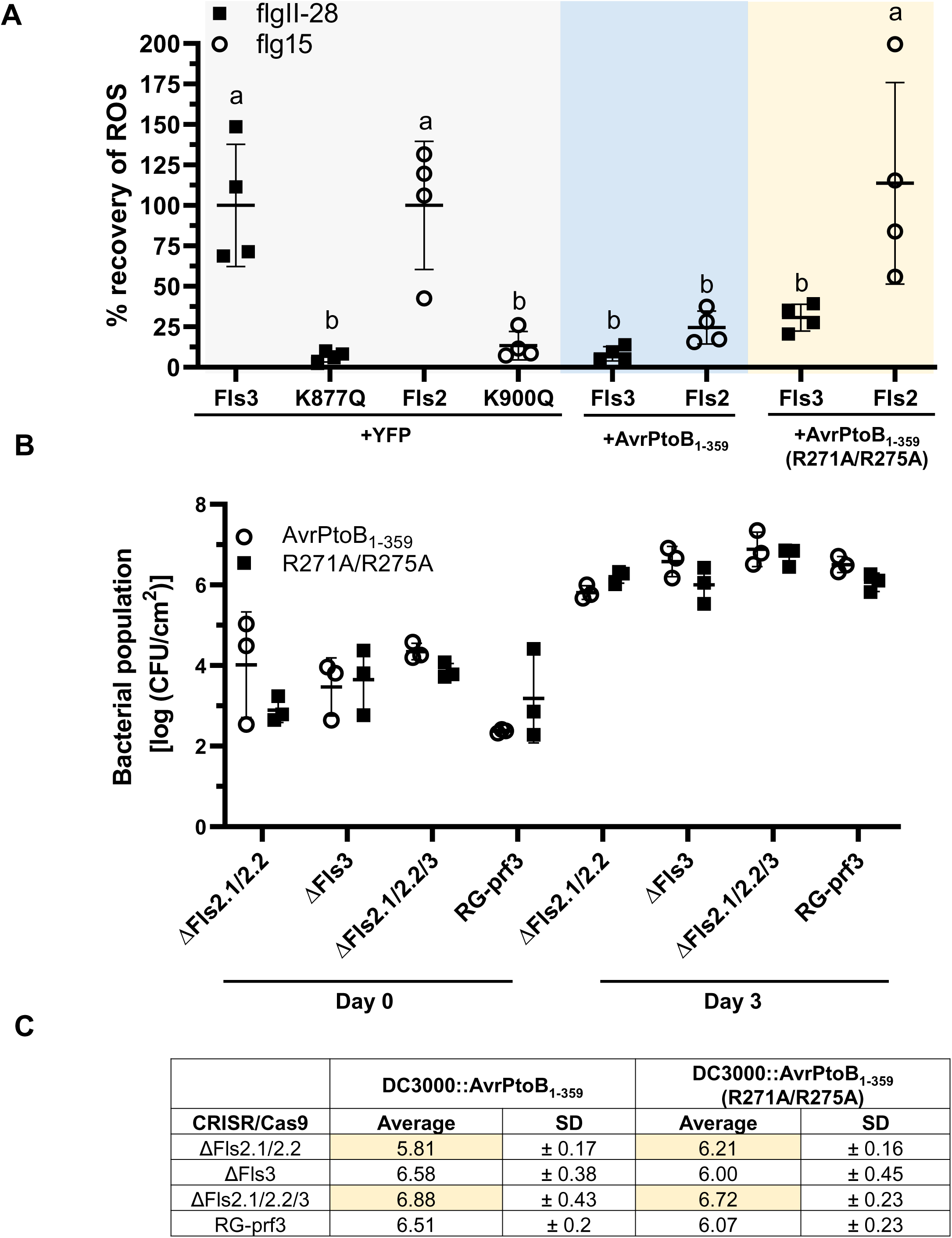
Fls3 and Fls2 are inhibited differently by the bacterial effector AvrPtoB. **(A)** Fls3 or Fls2 or their kinase inactive variants (K877Q or K900Q, respectively) were co-infiltrated with YFP or the AvrPtoB variants (AvrPtoB_1-359_ or AvrPtoB_1-359_ (R271A/R275A) into *N. benthamiana* leaves expressing a hairpin construct to knock down expression of *Prf* (hpPrf). Constructs were overexpressed under a 35S promoter. Samples were taken for ROS assays 48 hours post infiltration, and data shown are the means of four individual plants indicated as separate points, and horizontal lines are the means of the three plants +/- s.d.. Means are the total ROS values in response to 100 nM flg15 (Fls2) or flgII-28 (Fls3) peptide, accumulated over 45 minutes and normalized to the mean of Fls3 or Fls2 (100% ROS) to allow direct comparison of the Fls3 and Fls2 ROS values. Data are representative of four independent experiments. Statistical significance was determined using two-way ANOVA with Tukey’s multiple comparisons post-test with statistical significance cutoff *P*<0.05. **(B)** Bacterial populations in tomato leaves of mutants of *Fls2.1* and *Fls2.2* (ΔFls2.1/2.2), *Fls3* (ΔFls3), or *Fls2.1, Fls2.2*, and *Fls3* (ΔFls2.1/2.2/3) or Rio Grande-prf3 (RG-prf3) were measured 0 and 3 days after dip-inoculating plants with bacterial suspensions (1 x 10^8^ cfu/mL) using a DC3000 strain expressing AvrPtoB_1-359_ (labeled AvrPtoB_1-359_) or its variant AvrPtoB_1-359_(R271A/R275A) (labeled R271A/R275A). Shown are the means of three individual plants indicated as separate points, and horizontal lines are the means of the three plants +/- s.d.. There were no statistically significant differences using a pairwise t-test in bacterial growth between the two strains for each plant (*P*>0.05). **(C)** Table showing the averages and standard deviations (SD) of the data shown in **(B)** on Day 3. Data are from a single experiment and are representative of two replicates. *See also Figure S4*.

To further investigate whether AvrPtoB_1-359_(R271A/R275A) acts differently with Fls2 and Fls3, we tested if bacterial growth was affected. In RG-prf3, DC3000 harboring a plasmid encoding AvrPtoB_1-359_(R271A/R275A) (DC3000::AvrPtoB_1-359_(R271A/R275A)) grew slightly less than the wild type (DC3000::AvrPtoB_1-359_) in tomato leaves, though this difference was not statistically significant (Cheng et al., 2011). We vacuum infiltrated DC3000::AvrPtoB_1-359_ or DC3000::AvrPtoB_1-359_(R271A/R275A) into RG-prf3, ΔFls2.1/2.2, ΔFls3, or ΔFls2.1/2.2/3 plants and measured bacterial populations in leaves 3 days later to test whether Fls3 or Fls2 might interact differently with AvrPtoB when expressed by bacteria. In agreement with the previously published results in (Cheng et al., 2011), we observed a slight, but not statistically significant, reduction in growth of DC3000::AvrPtoB_1-359_(R271A/R275A) compared to DC3000::AvrPtoB1-359 in Rio Grande-prf3 (Fig. S4BC). However, in ΔFls2.1/2.2 plants the slight difference in growth between the DC3000::AvrPtoB_1-359_ and DC3000::AvrPtoB_1-359_(R271A/R275A) was lost, suggesting that PTI is no longer suppressed which allows the DC3000::AvrPtoB_1-359_(R271A/R275A) strain to grow more than in wild-type plants. In ΔFls3 plants, a similar difference in bacterial growth of DC3000::AvrPtoB_1-359_ and DC3000::AvrPtoB_1-359_(R271A/R275A) was observed as the wild type, suggesting that PTI is not suppressed and growth of DC3000::AvrPtoB_1-359_(R271A/R275A) is inhibited. In ΔFls2.1/2.2/3 plants, the slight difference in growth between the DC3000::AvrPtoB_1-359_ and DC3000::AvrPtoB_1-359_(R271A/R275A) was lost (Fig. S4BC).

Because PTI can have a greater effect on bacterial growth at the leaf surface, we dip inoculated the plants with DC3000::AvrPtoB_1-359_ or DC3000::AvrPtoB_1-359_(R271A/R275A) and measured bacterial growth in leaves to see if the difference in bacterial growth would be greater between the two strains. Although we still did not observe statistically significant differences in bacterial populations by pairwise t-tests in any of the Fls2 or Fls3 mutant or RG-prf3, we did observe a similar trend between the vacuum-infiltrated and dip-inoculated plants. In the dip-inoculated plants, there was a slight (but not statistically significant) reduction in the bacterial population of DC3000::AvrPtoB_1-359_(R271A/R275A) compared to wild-type DC3000::AvrPtoB_1-359_ in the RG-prf3 and ΔFls3 plants, but no reduction was observed in ΔFls2.1/2.2 or ΔFls2.1/2.2/3 plants. We confirmed protein expression of AvrPtoB_1-359_ and AvrPtoB_1-359_(R271A/R275A) in DC3000 via immunoblotting (Fig. S4D). Together, these data suggest that Fls3 and Fls2 interact differently with AvrPtoB *in planta*.

### Fls3 and Fls2 have different kinase activities *in vitro*

It is possible that different kinase activities of Fls3 and Fls2 account for their differences in immunity output. It was previously reported that Arabidopsis Fls2 has weak kinase activity *in vitro* that requires the entire intracellular domain for activity (both the inner juxtamembrane domain and kinase domain) (Cao et al., 2013; Gomez-Gomez et al., 2001; Lu et al., 2010; Xiang et al., 2008). To test whether tomato Fls3 and Fls2 have *in vitro* kinase activity and if their activity is dependent on the presence of the inner juxtamembrane domain (JM), we made recombinant proteins in BL21 *E. coli* that contained the Fls3 or Fls2 kinase domain (KD) alone, or the entire cytoplasmic domain containing the JM plus KD (JMKD), expressed in the pDEST-HisMBP vector (which has an N-terminal 6xHis-MBP tag), and assayed their kinase activity. As negative controls, we generated variants with substitutions in their ATP binding site, Fls2-JMKD(K900Q) and Fls3-JMKD(K877Q), and a control that expressed a short *E. coli* sequence to allow expression of the His-MBP protein in the pDEST-HisMBP vector. We found that both tomato Fls2 and Fls3 have kinase activity that depends on the presence of the inner juxtamembrane domain (Fig. 5A), and that the *in vitro* kinase activity for Fls3 is greater than Fls2 (Fig. 5). We added myelin basic protein (MyBP) to the *in vitro* kinase assays to test whether Fls2 and Fls3 could transphosphorylate this generic substrate. We observed that Fls3, but not Fls2, could transphosphorylate MyBP (Fig. 5B). Because Fls2 and Fls3 both require the inner juxtamembrane domain for *in vitro* kinase activity, we wanted to know if they required their own cognate inner juxtamembrane domain for activity. We swapped the JM domain between Fls3 and Fls2 (2JM-3KD and 3JM-2KD) and tested them in *in vitro* kinase assays. We found that swapping the inner juxtamembrane domains between Fls2 and Fls3 completely abolished *in vitro* kinase activity for both proteins (Fig. 5C). These data suggest that Fls3 and Fls2 have differences in their kinase activity that require their corresponding juxtamembrane domain.

**Figure 5.**
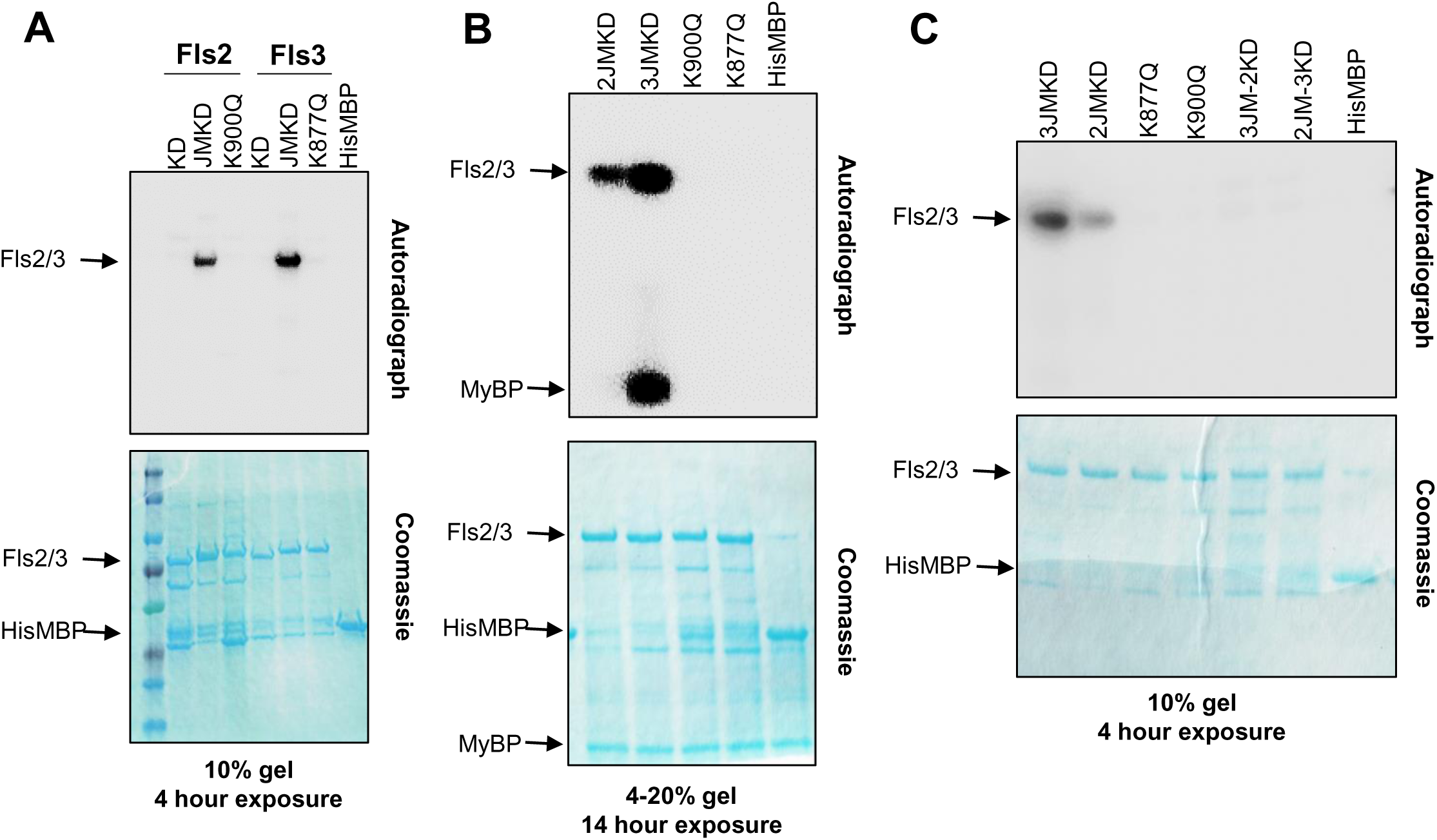
Fls3 and Fls2 vary in their kinase activity *in vitro*. The kinase domain alone (KD) or inner juxtamembrane domain plus the kinase domain (JMKD) of Fls3 and Fls2 were expressed as recombinant proteins in the pDEST-HisMBP vector and tested for *in vitro* kinase activity. The kinase inactive mutants for Fls3(K877Q) and Fls2(K900Q) were generated in the background of the JMKD constructs. HisMBP protein was used as a negative control. **(A)** Fls3 or Fls2 kinase domains (KD) or kinase domains plus the inner juxtamembrane domains (JMKD) were tested in *in vitro* kinase assays. **(B)** Myelin basic protein (MyBP) was added to the *in vitro* kinase assays to test the ability of the kinases to transphosphorylate the MyBP substrate. **(C)** Kinase assays swapping the inner juxtamembrane domains of Fls3 and Fls2 to test the requirement for their cognate inner juxtamembrane domain. 3JM-2KD expresses the inner juxtamembrane domain from Fls3 and the kinase domain from Fls2. 2JM-3KD expresses the inner juxtamembrane domain from Fls2 and the kinase domain from Fls3. Data shown are representative of at least three independent experiments. Below each panel is the exposure length to the phosphor-screen (4 or 14 hours) and the SDS-PAGE gel percentage.

### Subdomain I contributes to the *in vitro* differences in kinase activity for Fls3 and Fls2

We hypothesized that subdomain I of the kinase domain, which is involved in the correct positioning of the ATP substrate in the active site, contributes to the differences in kinase activity we observed between Fls2 and Fls3. We aligned the Fls2 (Fls2.1 and Fls2.2) and Fls3 subdomain I sequences with other immunity-associated tomato kinases and found that Fls2 deviates from the glycine-rich motif (GxGxxG) present in other kinases (Fig. 6A). Fls2 has the conserved, first position glycine, but the second and third glycines are replaced by serines (GxSxxS). Fls3 has the standard GxGxxG motif. We compared the sequence motif of all 42 tomato receptor-like kinases (RLKs) in the Fls2/Fls3 class (class IX-a) (Wei et al., 2015). We aligned the subdomain I kinase domain region of the 42 RLKs to determine the conservation and/or consensus of each of the residues in the GxGxxG motif. Two genes in this class, Solyc10g085110 and Solyc03g118330, likely do not encode active kinases because they lack the subdomain I region and the lysine responsible for binding of the ATP (Fig. S5A). We found that of the 40 RLKs in this class that have a subdomain I, only Fls2 deviates from the GxGxxG motif at the second position glycine (Fig. 6B and Fig. S5A). The third position glycine is more flexible in the class, with 28 RLKs harboring a glycine, 10 RLKs with a serine, 1 RLK with a valine, and 1 RLK with an isoleucine. The first position glycine is conserved in all 40 RLKs. To test whether the serines in the Fls2 subdomain I motif impact Fls2 kinase activity, we made substitutions of each serine individually or simultaneously to glycines (GxSxxS to GxGxxS (S881G), GxSxxG (S884G), or GxGxxG (S881G/S884G)). We found that the S881G mutation alone increased the kinase activity of Fls2 (Fig. 6C). In contrast, S884G resulted in a nearly complete loss of kinase activity. However, mutating both residues (S881G/S884G) led to an even stronger increase of kinase activity than S881G alone (Fig. 6C). To see if substituting the glycines to serines in the Fls3 subdomain I motif would impact kinase activity, we mutated the glycines to serines either individually or simultaneously (GxGxxG to GxSxxG (G858S), GxGxxS (G861S), or GxSxxS (G858S/G861S)). We found that the G858S mutation alone caused a near complete loss of *in vitro* kinase activity. The G861S mutation resulted in a strong reduction of kinase activity compared to the wildtype 3JMKD. Mutating both residues (G858S/G861S) also resulted in a reduction of kinase activity that was slightly less than G861S alone (Fig. 6C). These data support the hypothesis that the subdomain I motif contributes to the increased kinase activity of Fls3 compared to Fls2 *in vitro*.

**Figure 6.**
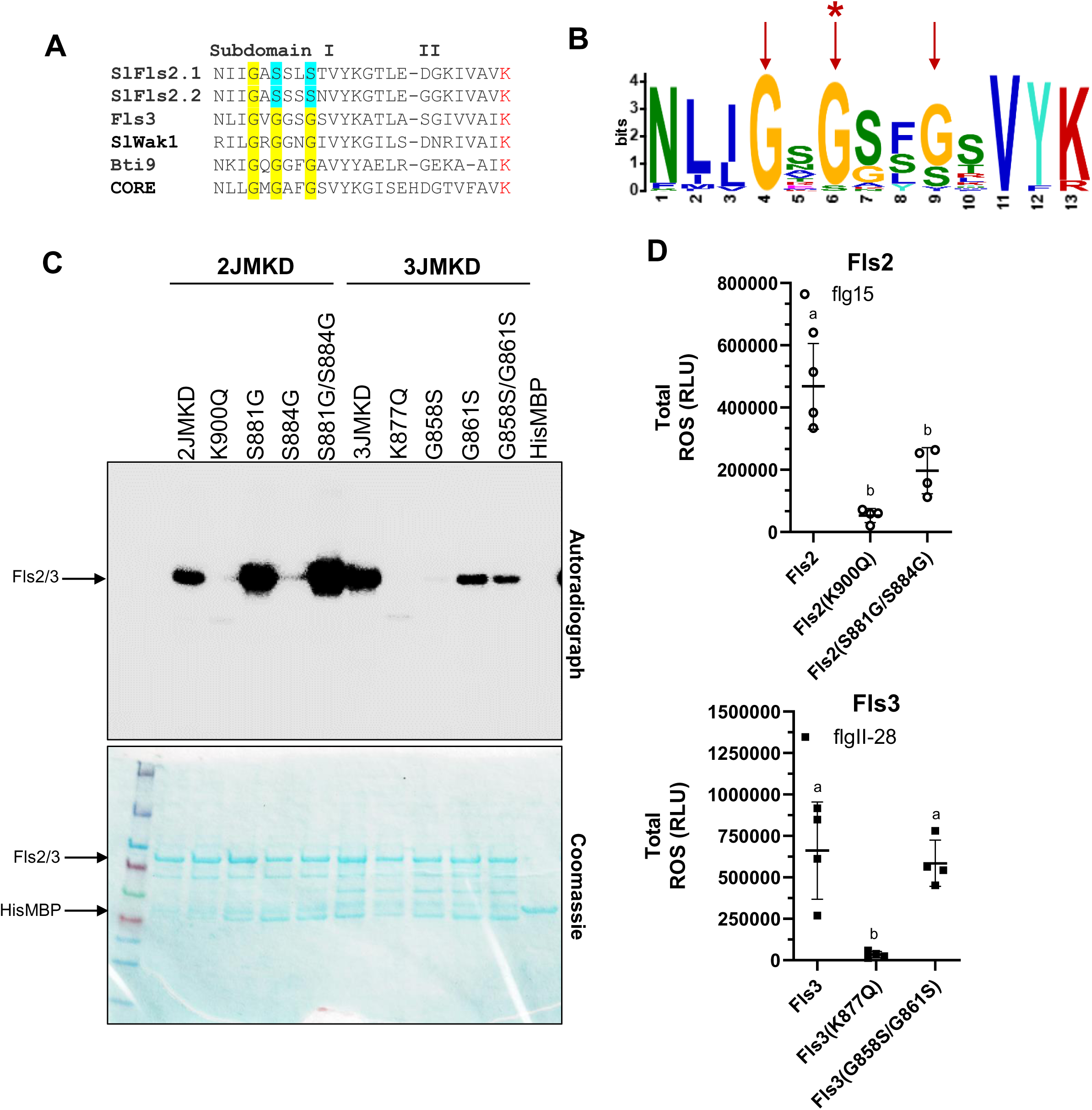
Subdomain I contributes to the differences in kinase activity between Fls3 and Fls2 *in vitro*. **(A)** Subdomain I in tomato showing the GxGxxG motif. In red is the lysine involved in ATP binding. Highlighted in yellow are the conserved glycines in the motif, and in blue are the deviant serines in Fls2. **(B)** MEME LOGO showing the prevalence of the GxGxxG motif in class IXa of the tomato RLKs, which includes Fls2 and Fls3. Red arrows indicate the three glycines in the GxGxxG motif. Only Fls2 has a different residue in the second position glycine (red asterisk). **(C)** *In vitro* kinase activity of Fls2JMKD and Fls3JMKD containing mutations in subdomain I. Data are representative of three independent replicates. **(D)** Total ROS accumulated over 45 minutes after applying 100 nM flg15 (Fls2 proteins) or flgII-28 peptide (Fls3 proteins) on leaf discs collected after transient expression of the various proteins in *N. benthamiana*. Protein overexpression was driven by the CaMV 35S promoter. Shown are the means of four individual plants indicated as separate points, and horizontal lines are the means of the four plants +/- s.d.. Data are from a single experiment and are representative of three independent replicates. Significance was determined via ANOVA followed by Tukey’s post-test with a significance cutoff of *P<*0.05. *See also Figure S5*.

We next tested whether substitutions in subdomain I of Fls3 and Fls2 affect the ROS output in transient *in planta* assays in the context of the full-length proteins. We cloned the double mutants Fls2(S881G/S885G) and Fls3(G858S/G861S) into the pGWB417 plant protein expression vector that has a C-terminal myc-tagged protein, cloned the kinase inactive pGWB417::Fls2(K900Q) and pGWB417::Fls3(K877Q) as negative controls, and transiently expressed the proteins in *N. benthamiana* leaves. ROS responses to 100 nM flgII-28 or flg15 peptide were measured and compared to the wild-type Fls2 or Fls3 proteins. Surprisingly, we found that there was a statistically significant reduction in total ROS production of Fls2(S881G/S885G) compared to wildtype Fls2. There was no statistically significant difference in total ROS production of Fls3(G858S/G861S) compared to wildtype Fls3. We confirmed that all proteins were transiently expressed via immunoblotting (Fig. S5B). These results suggest that although the subdomain I sequence influences *in vitro* kinase activity, the differences in kinase activity alone do not explain the differences in ROS output between Fls2 and Fls3.

### No single domain explains the sustained ROS response of Fls3

It is also possible that differences in their leucine rich repeat (LRR) domains, which are the extracellular domains that bind the flg22 or flgII-28 peptides, are responsible for the different readouts of these two PRRs. To test this hypothesis, we generated chimeric constructs that swapped the LRR, transmembrane (TM), and kinase domains (KD) between Fls2 and Fls3 for transient expression in *N. benthamiana*. We included the presumed signal peptide and N-terminus within the ‘LRR’ domain, the inner- and outer-juxtamembrane domains within the ‘TM’ region, and the C-terminus within the ‘KD’ (Fig. 7A). We used a low concentration of peptide in the ROS assays (10nM flgII-28 or flg15) to allow detection of subtle differences in the ROS responses of the chimeric constructs. As a control, we also generated the reconstituted wild-type constructs, which have the same sequence as the wild-type Fls3 or Fls2 constructs but were generated in the same manner as the chimeric constructs (labeled 3-3-3 or 2-2-2, respectively).

**Figure 7.**
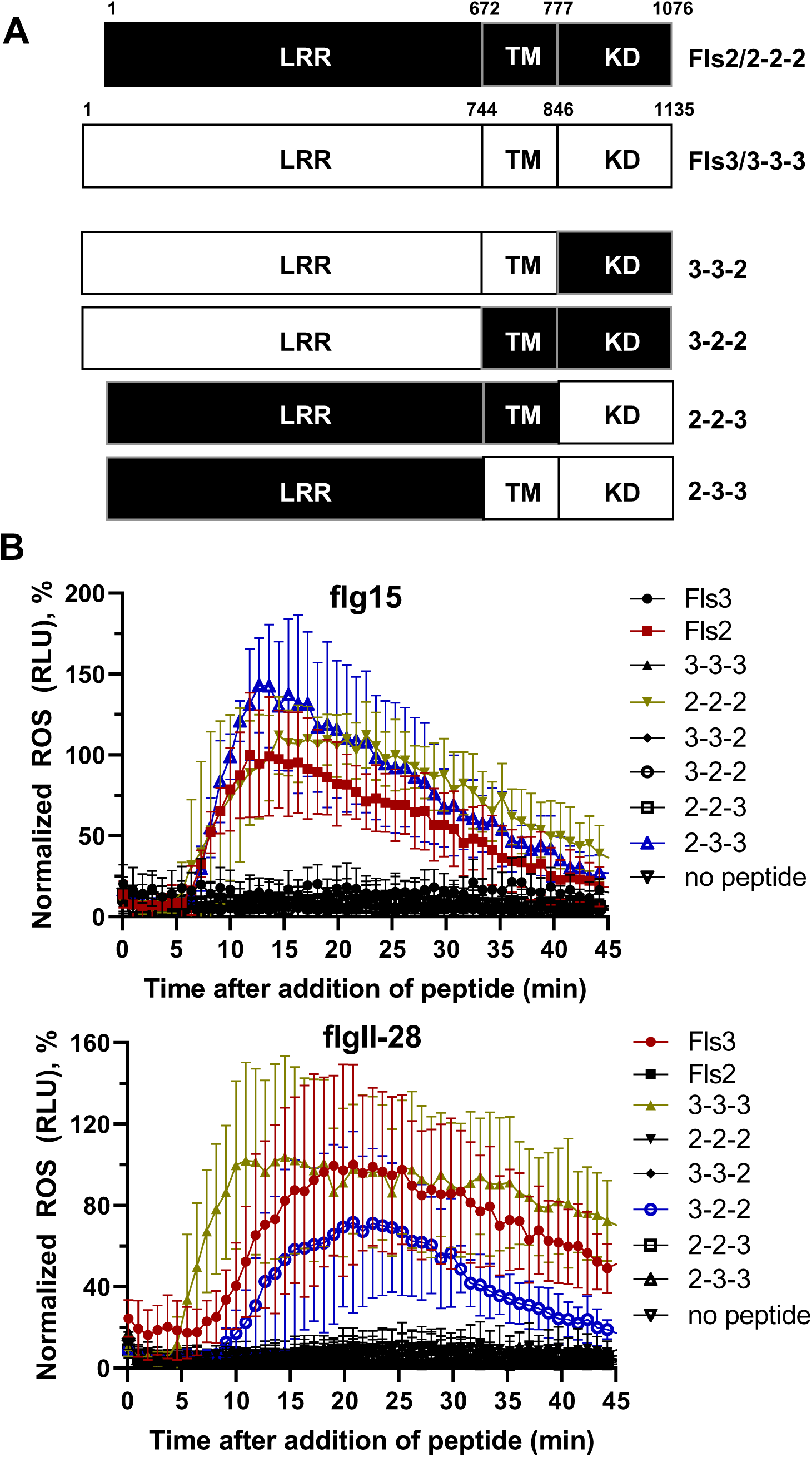
No single domain governs the sustained ROS response. **(A)** Schematic diagram showing the design of the chimeric Fls2 and Fls3 constructs. Amino acid numbers corresponding to the domains used to generate the chimeric proteins are shown above each schematic. Leucine rich repeat, LRR; transmembrane domain, TM; and kinase domain, KD. The labels for each chimeric construct are as follows: the ‘2’ or ‘3’ indicates whether the domain came from Fls2 (‘2’) or Fls3 (‘3’). The first number indicates the LRR, second number indicates the TM, and third number indicates the KD. The reconstituted Fls2 (2-2-2) and Fls3 (3-3-3) were included as controls and are identical in sequence to wildtype Fls2 or Fls3, respectively. **(B)** ROS curves comparing the chimeric constructs for the sustained response. Constructs were transiently expressed in *N. benthamiana* leaves under control of the CaMV 35S promoter and ROS activity was measured over 45 minutes after the addition of 10 nM flg15 or flgII-28 peptide. Data are the means of four plants (n=4), and error bars represent +/-s.d. and are representative of three independent experiments. The two chimeric constructs that responded to peptide are highlighted in blue. The unadulterated controls are highlighted in red, and the reconstituted controls are highlighted in gold. *See also Figure S6*.

In our transient assays, we found that only the chimeric constructs with the TM and KD from the same receptor responded to peptide (2-3-3 and 3-2-2; Fig. 7B). Specifically, the chimeric construct with the LRR from Fls2 and the TM and KD from Fls3 (2-3-3) responded similarly as the wildtype Fls2 and reconstituted wildtype construct (2-2-2) to the flg15 peptide, both in amplitude and duration of response (Fig. 7B and S6AB). The time at which the maximum ROS response to flg15 peptide occurred was also similar between 2-3-3, Fls2, and 2-2-2 (about 12 minutes after peptide treatment). The normalized ROS responses at the maximum amplitudes were not statistically different between 2-3-3, Fls2, and 2-2-2 (Fig. S6A). Additionally, 45 minutes after peptide treatment there was no significant difference between the ROS responses of 2-3-3, Fls2, and 2-2-2 (Fig. S6B).

When we applied 10 nM of the flgII-28 peptide, only the chimeric construct with the LRR from Fls3 and the TM and KD from Fls2 (3-2-2) activated a ROS response (Fig. 7B). However, while the time at which the maximum response occurred was similar between 3-2-2, wildtype Fls3, and the reconstituted wildtype construct (3-3-3) (about 20 minutes after peptide treatment) and the normalized maximum amplitudes of the ROS responses were not statistically significant (Fig. 7B and Fig. S6A), the ROS response for 3-2-2 was not sustained like Fls3 and 3-3-3 (Fig. 7B and Fig. S6B). At 45 minutes after flgII-28 treatment, the ROS response for 3-2-2 was significantly less than Fls3 and 3-3-3, and was not significantly different from the flgII-28 non-responders (Fig. S6B). All proteins were confirmed to be expressed via immunoblotting, though there were some differences in the levels of expression between the chimeric constructs (Fig. S6C). However, expression of 3-2-2 was similar to Fls3 and 3-3-3, so expression alone does not explain the differences in the sustained ROS responses between Fls3 and 3-3-3 and 3-2-2. Additionally, the expression of 2-3-3 was reduced compared to Fls2 and 2-2-2 (to levels similar to Fls3 and 3-3-3) but 2-3-3 responded similarly to peptide as Fls2 and 2-2-2 and this therefore supports that expression levels alone do not account for the differences observed between the chimeric constructs. The data in Fig. 7 and Fig. S6 were normalized to the in-plate Fls3 or Fls2 controls to also account for differences in expression between constructs, plants, and technical replicates. Together, these data suggest that no single domain is responsible for the sustained ROS response.

## Discussion

While both Fls2 and Fls3 recognize flagellin-derived MAMPs, their individual contributions to disease resistance was unknown. We used the CRISPR/Cas9 system to generate mutations in the *Fls3* and two *Fls2* genes in tomato and used them to determine the contributions of the two PRRs to disease resistance. Both Fls3 and Fls2.1 were found to have similar effects on inhibition of *Pst* growth on the leaf surface and in the leaf apoplast, with the greatest effects occurring on the leaf surface (Fig. 1 and S1). Additionally, we found that a 7 base pair deletion in Fls2.1 alone completely abolished flg22 recognition in tomato, further supporting the idea that Fls2.1 is the only functional receptor for flg22 recognition in tomato leaves as reported previously (Jacobs et al., 2017).

One observation that suggested there may be differences in the molecular mechanisms between Fls2 and Fls3 was that the ROS response to flgII-28 is sustained compared to flg22 (Fig. 2 and S2). We observed this sustained ROS response in both tomato and potato, and data from another study appears to support a sustained ROS response for flgII-28 in pepper (Clarke et al., 2013). While others have reported differences in the amplitude of flgII-28 and flg22 ROS responses in potato (Moroz and Tanaka, 2019), we did not observe these differences under our conditions. This may be attributed to the use of different potato varieties or peptide concentrations in the assays (100nM in our study versus 1μM in (Moroz and Tanaka, 2019)). Additionally, although *N. benthamiana* does not have an endogenous copy of *Fls3*, transient overexpression of Fls3 in *N. benthamiana* also results in a sustained response compared to Fls2 (this study and (Hind et al., 2016)). Previous efforts to express Fls3 in Arabidopsis have been unsuccessful (Hind et al., 2016), so it is currently unknown whether Fls3 can function in species outside the *Solanaceae,* but future efforts to express Fls3 in more diverse species could determine whether Solanaceous-specific protein partners are involved in the Fls3 ROS response.

We analyzed RNA-Seq data from previous studies (Pombo et al., 2017; Rosli et al., 2013) and found four genes that were specifically induced upon application of flgII-28 peptide (Fig. 3A). We measured transcript abundance of two of these genes, *BHLH* and *UGT*, six hours after treatment of leaves with flgII-28 or flg22 peptide and found that both genes were specifically induced by flgII-28 induction (Fig. 3B). This suggests these two genes are involved in an Fls3-specific signaling pathway that is independent of Fls2 and supports the hypothesis that there are differences in the molecular signaling pathways between Fls3 and Fls2. We previously identified three genes whose expression is induced specifically during PTI (Pombo et al., 2014). These genes were not identified as specific Fls3 reporters in the RNA-Seq data indicating they may be induced by both flg22 and flgII-28. While we have yet to find an Fls2-specific reporter gene, future studies using pathway-specific reporter genes may help dissect the differences in the signaling components activated by Fls2 and Fls3.

AvrPtoB inhibits Fls2 by binding to BAK1, and possibly also by directly binding to Fls2 (Cheng et al., 2011; Gohre et al., 2008; Rosebrock et al., 2007; Shan et al., 2008). The R271 and R275 residues in AvrPtoB are important for binding to BAK1, and mutating these residues results in a loss of interaction with BAK1 and a slight reduction in bacterial growth due to the failure of AvrPtoB to suppress PTI (although this reduction is not statistically significant) (Fig. 4 and (Cheng et al., 2011)). We transiently co-expressed Fls2 or Fls3 and AvrPtoB_1-359_ or its variant AvrPtoB_1-359_(R271A/R275A) in *N. benthamiana* leaves and found that both Fls2 and Fls3 ROS responses were inhibited upon co-infiltration with AvrPtoB_1-359_ (Fig. 4A). However, when Fls2 and Fls3 were co-infiltrated with AvrPtoB_1-359_(R271A/R275A), this AvrPtoB variant still suppressed Fls3 although it did not suppress the Fls2 ROS response and Fls2-driven PTI. These data suggest there may be differences in the molecular interaction between Fls2 and Fls3 and AvrPtoB. Additionally, because AvrPtoB interacts with BAK1 it is possible that Fls2 and Fls3 interact differently with the tomato homologs of BAK1 (SERK3A/SERK3B) and this will be a focus of future studies.

It has been previously reported that Arabidopsis Fls2 has weak autophosphorylation activity *in vitro* and *in vivo* that requires the presence of the inner juxtamembrane domain (Cao et al., 2013; Gomez-Gomez et al., 2001; Lu et al., 2010; Xiang et al., 2008). We found that tomato Fls2 and Fls3 both have relatively strong autophosphorylation activity *in vitro* that can be detected after only four hours of exposure to a phosphor-screen. This activity is dependent on the presence of the inner juxtamembrane domain, and it is possible that this may be due to specific important residues within the domain. For example, a previous study examined the requirement of the inner juxtamembrane domain for the rice PRR Xa21 function, and found that the C-terminal region of the inner juxtamembrane domain was required for autophosphorylation (Chen et al., 2010). The authors found a conserved threonine residue (T705) within the juxtamembrane domain at the C-terminal end of the domain that was conserved among plant RLKs, and mutation of this residue (T705A or T705E) resulted in increased susceptibility to *Xanthomonas oryzae* pv. oryzae. It is still unknown what molecular role this residue and the inner juxtamembrane domain as a whole play in kinase activity, but the authors propose that the threonine may be serving a dual role in receiving and donating a phosphoryl group (Chen et al., 2010). Tomato Fls2 and Fls3 both have this conserved threonine residue, and future research is needed to determine the molecular role of their inner juxtamembrane domains. Some other plant RLK chimeras have been shown to require the cognate inner juxtamembrane domain for their function. For example, a chimeric construct containing the extracellular domain of the rice chitin elicitor receptor CEBiP and the kinase domain from the rice blast resistance protein Pi-d2 was only functional if the transmembrane domain originated from Pi-d2 (Kouzai et al., 2013). However, not all RLKs require their cognate inner juxtamembrane domain to function. Swapping the inner juxtamembrane domains of the Arabidopsis RLK CERK1 with BAK1 and Fls2 still resulted in a functional CERK1 protein (Zhou et al., 2019). Our future efforts will focus on understanding why there is a cognate inner juxtamembrane requirement for Fls2 and Fls3.

Subdomain I of the Fls3 kinase domain contributes to the stronger *in vitro* kinase activity of this protein (Fig. 5, 6). A previous study speculated that substituting the second glycine for a serine, as seen in Arabidopsis FLS2 (S879), would lead to a major reduction in kinase activity for FLS2 and supports the data that Arabidopsis has weak autophosphorylation ability (Schwessinger et al., 2011). Therefore, one would predict that changing of S881 to S881G in tomato Fls2 would result in a dramatic increase in kinase activity (Fig. 6C). In fact, when we made this substitution we did observe a dramatic increase in kinase activity. However, the S884G substitution resulted in a complete abolishment of kinase activity, yet mutation of both residues (S881G/S884G) caused an even greater increase in kinase activity than S881G alone. Conversely, the G858S substitution in Fls3 caused a complete abolishment of kinase activity, and the G861S and G858S/G861S mutations resulted in a dramatic reduction, but not abolition, of Fls3 kinase activity (Fig. 6C). Further studies are needed to determine why Fls2 requires a serine at residue 884 in the context of GxSxxS or GxGxxS for kinase activity, but these differences between Fls3 and Fls2 suggest there may be differences in the molecular mechanisms of kinase activation between Fls2 and Fls3. Kinase activity alone, however, does not explain the sustained ROS response *in planta*. When we made the Fls2(S881G/S884G) and Fls3(G858S/G861S) mutations in the context of the full-length protein and overexpressed them in *N. benthamiana* leaves, the Fls2(S881G/S884G) mutation resulted in a decrease of total ROS production, rather than an increase. For the Fls3(G858S/G861S) mutation, we observed no statistically significant effect of the mutation on total ROS production. It is currently unknown why the *in vitro* results do not translate to the *in planta* ROS assays, but future experiments studying the kinase activity *in vivo* will help uncover the mechanisms of kinase activation in Fls3 and Fls2.

We hypothesized that either the kinase domain or the LRR domain may be responsible for the sustained ROS response observed for Fls3. Therefore, we generated chimeric Fls2 and Fls3 constructs to test whether either of these domains was solely responsible for the sustained ROS response (Fig. 7 and S6). When we overexpressed the chimeric constructs in *N. benthamiana* leaves and measured the ROS responses to flgII-28 or flg15, we discovered that there is a requirement for the transmembrane domain (TM) and the kinase domain (KD) to be from the same receptor, as only the constructs with a TM and KD originating from the same receptor responded to peptide (Fig. 7 and S6). This observation agrees with the *in vitro* kinase results that show that kinase activity requires the cognate inner juxtamembrane domain, and suggests that the lack of ROS responses in the chimeric constructs with a TM and KD from different receptors may be due to a lack in kinase activity (Fig. 5C). We also found that neither of the two chimeric constructs that responded to peptide (2-3-3 to flg15 or 3-2-2 to flgII-28) had a sustained ROS response (Fig. 7B and S6AB). While the shape of the curve and time at maximum amplitude resembled the receptor matching their LRR domain (12 min for Fls2 and 2-3-3, and 20 min for Fls3 and 3-2-2), both chimeric constructs were statistically indistinguishable from their negative controls at 45 minutes after peptide treatment, while Fls3 and 3-3-3 maintained the ROS levels at about 50-60% of their maximum amplitude (Fig. 7 and S6).

Collectively, our data suggest that no single domain is responsible for the sustained ROS response and that multiple structural features unique to Fls3 might contribute to the sustained response. While previous data show that Fls3 has some similar molecular characteristics as Fls2 (Hind et al., 2016), our data suggest that Fls3 acts in a different signaling pathway compared to Fls2 and interacts differently with the bacterial effector AvrPtoB. Determining the different components of the Fls3 and Fls2 pathways and how they differentially interact with *Pst* effectors may shed light on how Fls3 evolved as a solanaceous-specific flagellin receptor and help us better understand the molecular mechanisms of plant immunity.

## Methods

### Plant growth conditions, inoculations, and bacterial growth assays

Tomato seedlings were grown under the conditions described previously and inoculated as described in (Roberts et al., 2019b) (see the Supplemental Methods for more information). For the dip inoculations, three-week-old seedlings were placed in a 100% relative humidity chamber for 14 h prior to inoculation, then dipped into the bacterial suspension of 1 x 10^8^ CFU ml^-1^ for 10 seconds. Bacterial populations were quantified on Day 0 and Day 2 or 3 as described previously (Roberts et al., 2019b). See the Supplemental Methods section for more information. For a list of bacterial strains used in this study, see Table S5.

### Generation of CRISPR Cas9-mediated knockout lines

gRNAs were designed to target *Fls3*, *Fls2.1*, or both *Fls2.1*/*Fls2.2* as described previously (Jacobs et al., 2017; Zhang et al., 2020) using the tomato genome version SL2.5 (Tomato Genome Consortium, 2012). To induce mutations in both *Fls3* and *Fls2.1/2.2* in the same plant (ΔFls2.1/2.2/3), the constructs used to induce the individual mutations were transformed into the *Agrobacterium tumefaciens* strain LBA4404 and the cultures were mixed 1:1 prior to tomato transformation. Tomato transformations were performed at the Boyce Thompson Institute transformation facility (Gupta and Van Eck, 2016; Van Eck et al., 2019). More information may be found in the Supplemental Methods, Table S1, Table S4–S5, Fig. S1, and on the Plant CRISPR database at plantcrispr.org.

### Reactive oxygen species bioassays

ROS production was measured in response to flgII-28, flg22, or flg15 peptides as described in (Roberts et al., 2019b) using 100nM concentrations for all assays except for the chimeric constructs (10nM flgII-28 and flg15, Fig. 7 and Fig. S6). For Fig. 4 (effector suppression), total ROS accumulation was summed over 45 minutes and the values were normalized to either Fls3 or Fls2 so that all constructs could be directly compared (the samples did not all fit on a single plate). Every experiment was repeated at least three times with four plants per experiment and values represent the means of the four plants, +/-s.d as determined using the Prism 8 program. Shown is one representative replicate for each experiment. See the Supplemental Methods for information about the peptides used in this study.

### Reporter genes

Three, five-week-old Rio Grande-prf3 plants were syringe-infiltrated with 1μM flgII-28 or flg22 peptide or water and sampled 6 h post infiltration. Biological replicates were taken from each of the three plants infiltrated with peptide or water. RNA extraction, cDNA synthesis, and qRT-PCR was performed as described previously (Pombo et al., 2014), and significance was determined using a pairwise t-test using the Prism 8 program. Primers used for qRT-PCR may be found in Table S4. Gene ontology terms (GO Terms) were determined using the Plant Transcriptional Regulatory Map software (http://plantregmap.cbi.pku.edu.cn/go.php).

### Cloning

The chimeric constructs were generated via overlap extension PCR. The 2KD, 2JMKD, PCR products were inserted into the entry vector pJLSMART (Mathieu et al., 2014). Chimeric construct ORFs were then recombined into the Gateway vector pGWB417 (Nakagawa et al., 2009; Nakagawa et al., 2007) using the LR Clonase II following the manufacturer’s instructions (Thermo Fisher Scientific, https://www.thermofisher.com/us/en/home.html). Mutagenesis of the 2JMKD, 3JMKD, Fls2, and Fls3 clones was performed in the entry vectors using the Q5 Site-Directed Mutagenesis kit following the manufacturer’s instructions (New England Biolabs, www.neb.com). For a list of all primers and constructs used in this study, see Table S4 and S5. See the Supplementary Methods for more information.

### Agroinfiltrations

Agroinfiltrations of binary vectors into *N. benthamiana* were performed as previously described (Hind et al., 2016). All cultures were prepared to a final OD_600_ of 0.2. The Fls3- and Fls2-containing bacterial cultures were mixed 1:1 with a construct expressing the *p19* viral suppressor of silencing. See the Supplemental Methods for more information.

### *In vitro* kinase assays

HisMBP-tagged proteins were transformed into BL21 (DE3) pLys Rosetta cells and grown in culture at 37°C until the OD_600_ reached 0.6-0.8. Protein expression was induced using 1mM IPTG at 28°C for 3-4 h. Cell pellets were suspended in column buffer (20 mM Tris-HCl, pH 7.5, 200 mM NaCl, 1 mM EDTA, 1 mM DTT, 10% glycerol) supplemented with Complete Easy protease inhibitor cocktail (Millipore Sigma, https://www.sigmaaldrich.com/united-states.html), lysed by sonication, mixed with amylose resin (New England BioLabs, https://www.neb.com/), and eluted with 10 mM maltose. *In vitro* kinase assays were performed using 5 μg of each of the various kinase proteins and/or 3 μg of myelin basic proteins and conducted as described previously (Roberts et al., 2019a).

### Immunoblotting

For the transiently-expressed proteins in *N. benthamiana,* total protein was extracted from Agroinfiltrated leaves, 5-10 μg was run on SDS-PAGE, blotted on PVDF membrane, and detected with anti-Myc antibodies (Genscript, #A00704 www.genscript.com) and ECL Plus chemiluminescent substrate (Thermo Fisher Scientific, www.thermofisher.com) as described in (Roberts et al., 2019a).

To confirm the expression of AvrPtoB_1-359_ and AvrPtoB_1-359_(R271A/R275A) in DC3000 (DC3000::AvrPtoB_1-359_ and DC3000::AvrPtoB_1-359_(R271A/R275A), Fig. 4BC and Fig. S4), the hrp promoter on plasmids carrying AvrPtoB_1-359_ or AvrPtoB_1-359_(R271A/R275A) was induced as described previously (Cheng et al., 2011). Protein was equally loaded on SDS-PAGE for immunoblotting and the HA tag was detected using an anti-HA antibody (Roche, #11867423001 www.sigmaaldrich.com) and Femto chemiluminescent substrate (Thermo Fisher Scientific, www.thermofisher.com).

## Author Contributions

Conceptualization, R.R. and G.B.M; Methodology, R.R. and G.B.M; Investigation, R.R., A.E.L, L.W., A.M.G., S.R.H, and H.G.R.; Writing-Original Draft, R.R. and G.B.M; Writing-Review and Editing, R.R. and G.B.M.; Funding Acquisition, G.B.M.

## Acknowledgments

We thank Ning Zhang for helpful comments on the manuscript, Jing Zhang, Fabian Giska, and Ning Zhang for performing supporting experiments, Kevin Chen and Ben Carter for experimental assistance, Brian Bell, Jay Miller, and Nick Vail for greenhouse assistance, and Jason Ingram for providing potato tubers used in this study. Funding was provided by National Science Foundation grant IOS-1546625 (GBM). The authors declare no conflicts of interest.

---

**Figure S1.**
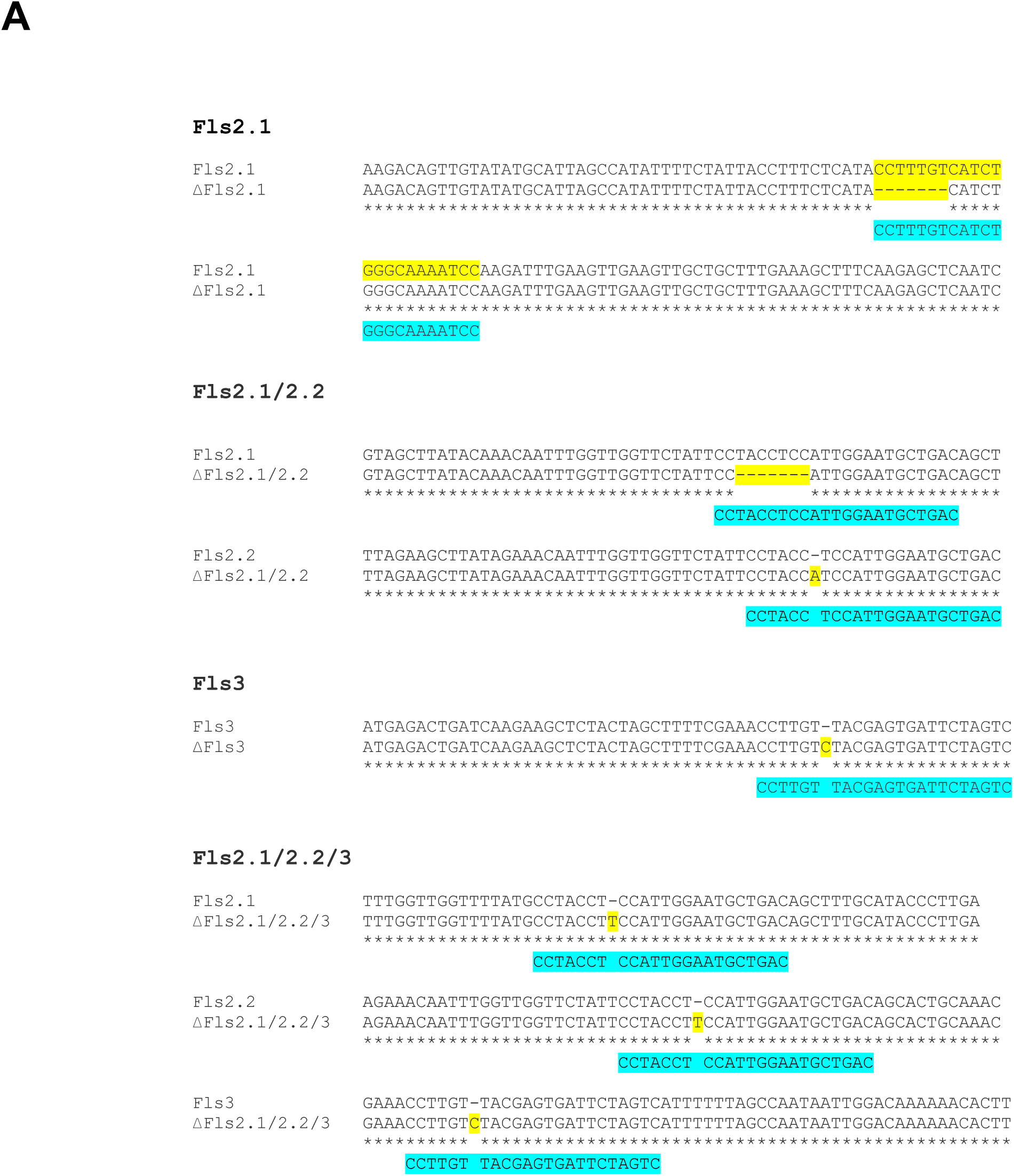
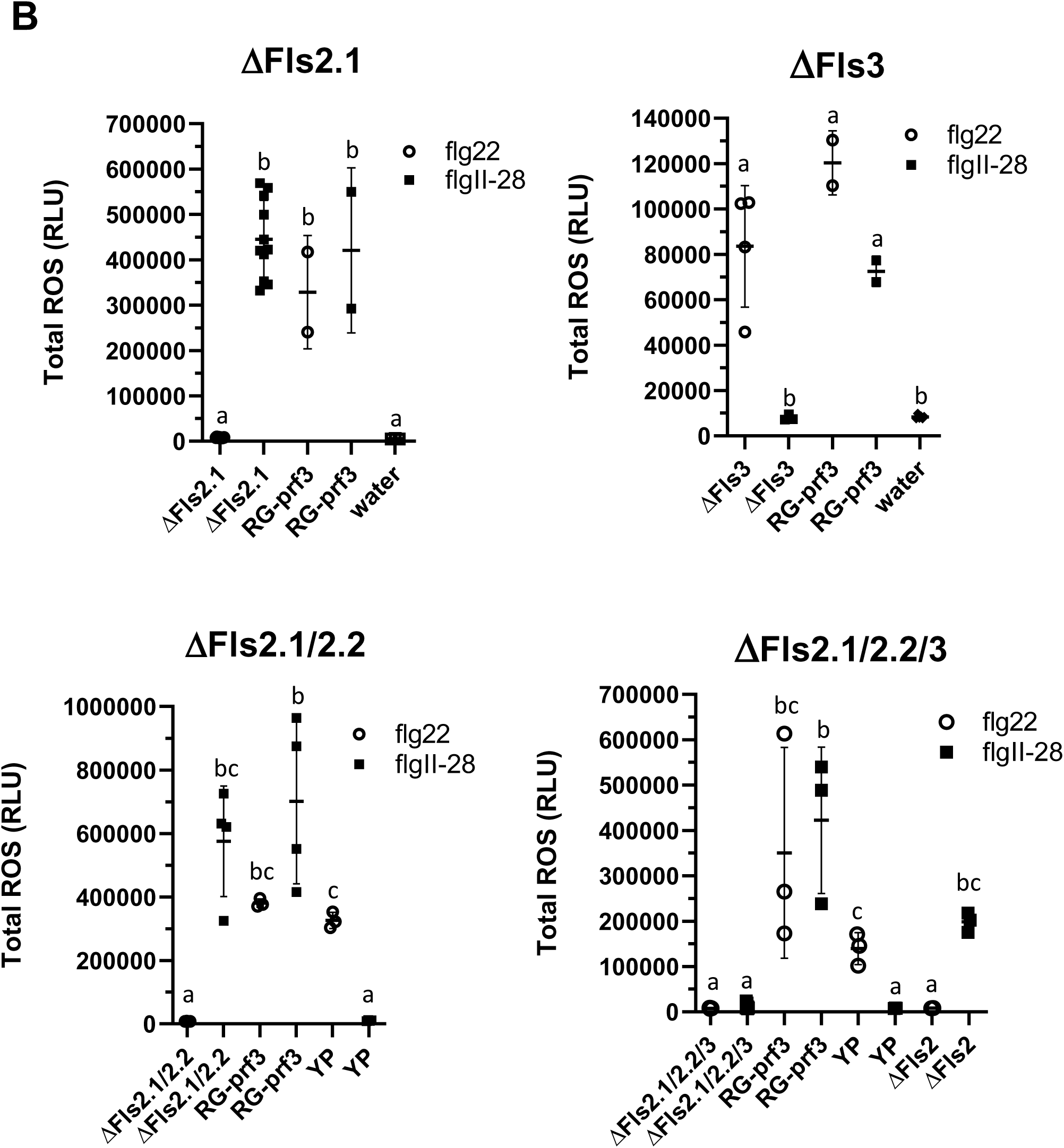
Development and characterization of CRISPR/Cas9-generated mutations in Fls2.1, Fls2.2 and Fls3. Related to Figure 1. **(A)** Alignment of CRISPR/Cas9-induced mutations (yellow) and gRNA target (blue) against each gene sequence. **((B),** next page) **(B)** ROS showing loss of flg22 and flgII-28 perception in Rio Grande-prf3 tomato plants with mutations in *Fls2.1* (ΔFls2.1), *Fls3* (ΔFls3), *Fls2.1* and *Fls2.2* (ΔFls2.1/2.2), or *Fls2.1, Fls2.2*, and *Fls3* (ΔFls2.1/2.2/3). Shown are ROS produced in response to 100nM flgII-28 or flg22 peptide or water in the mutant plants (in the background of Rio Grande-prf3), Rio Grande-prf3, Yellow Pear, or ΔFls2 in the background of M82 sp+ (Jacobs et al 2017). Results shown are the means of individual plants indicated as separate points, and horizontal lines are the means of the plants +/-s.d. (n=11 T1 plants for ΔFls2.1; n= 4 T2 plants for ΔFls3 and ΔFls2.1/2.2; n=13 T3 plants for ΔFls2.1/2.2/3; n=2 for Rio Grande, Yellow Pear, and ΔFls2). Significance was determined via one-way ANOVA with Tukey’s multiple comparisons test.

**Figure S2.**
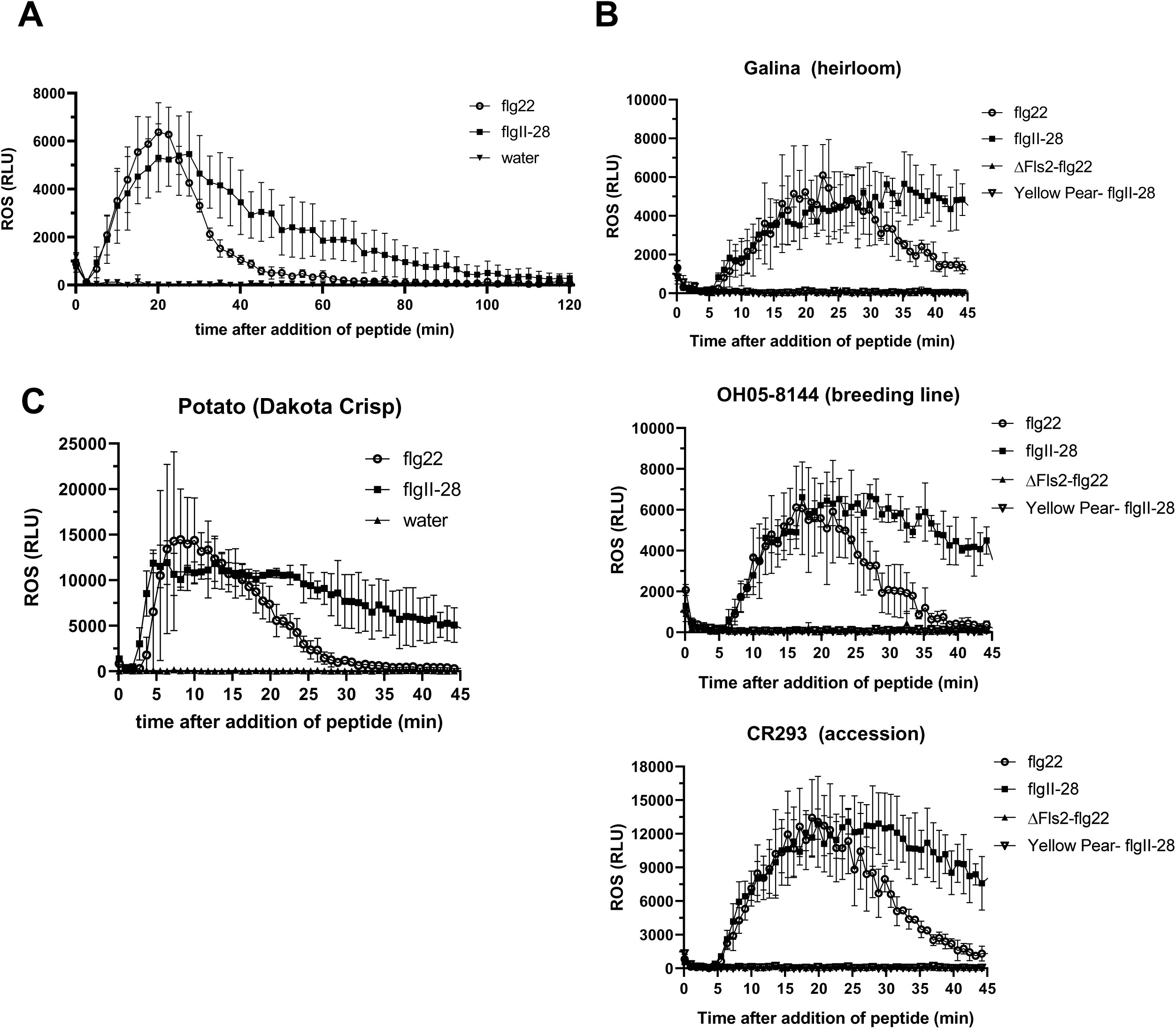
Additional tomato and potato accessions with sustained flgII-28 ROS response. Related to Figure 2. Oxidative burst (ROS) produced in response to 100nM flgII-28 or flg22 peptide or water in tomato and potato. **(A)** The tomato Rio Grande-prf3 flgII-28 response is sustained for 100 minutes, compared to 60 min for flg22. **(B)** Tomato heirloom Galina, breeding line OH05-8144, and accession CR293, or **(C)** Potato var. Dakota Crisp, all have sustained ROS responses to flgII-28. ROS was measured in relative light units (RLU). Results shown are means +/-s.d. (n=3 for tomato, n=2 for potato) and are representative of two (tomato) or three (potato) independent experiments.

**Figure S3.**
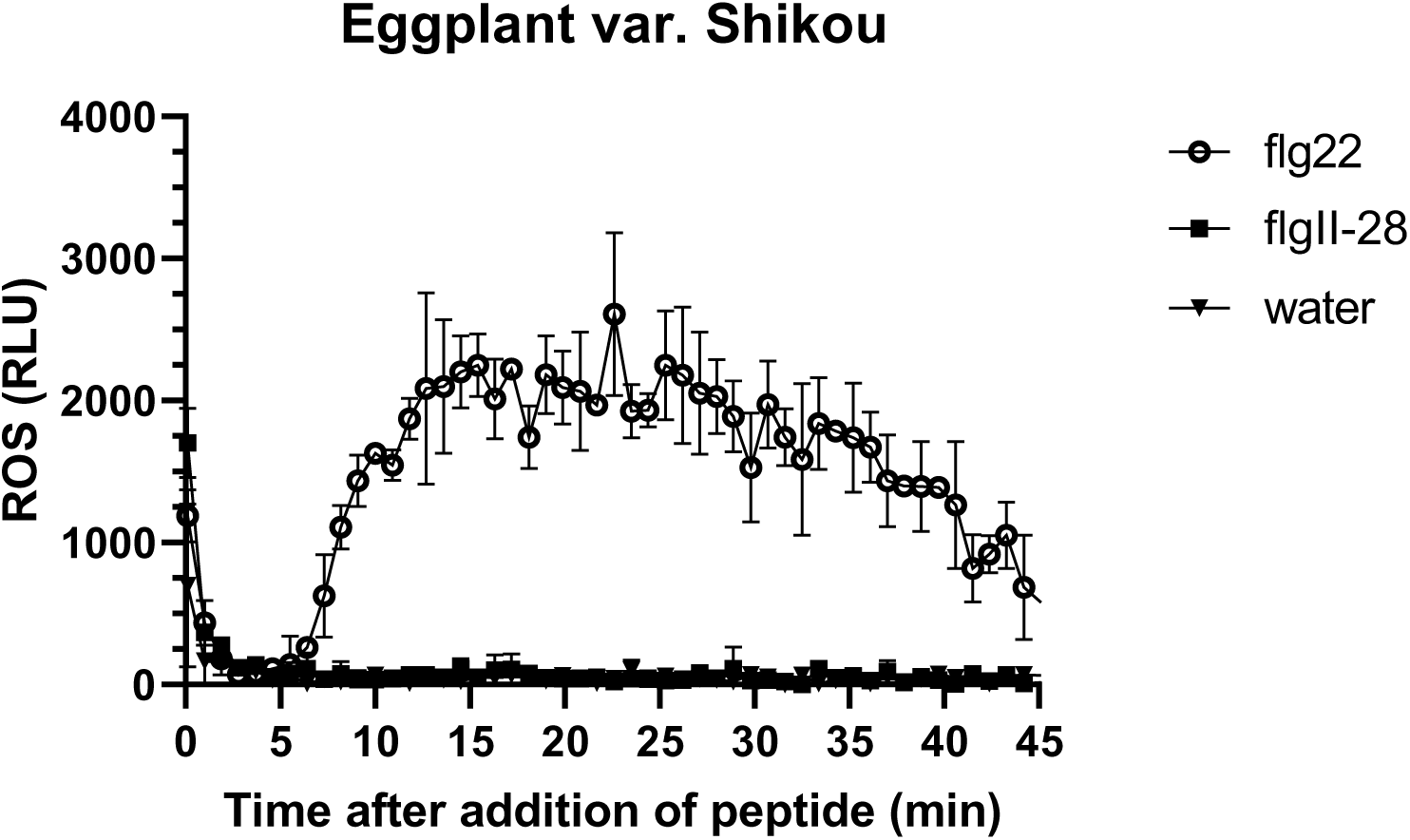
Eggplant var. Shikou does not respond to flgII-28. Related to Figure 2. Oxidative burst (ROS) produced in response to 100nM flgII-28 or flg22 peptide or water, measured in relative light units (RLU). Results shown are means +/-s.d. (n=3 plants). The four oldest leaves from each of three 6-week-old plants were tested, and shown are the results from leaf 4. Similar observations were made for each of the four leaves.

**Figure S4.**
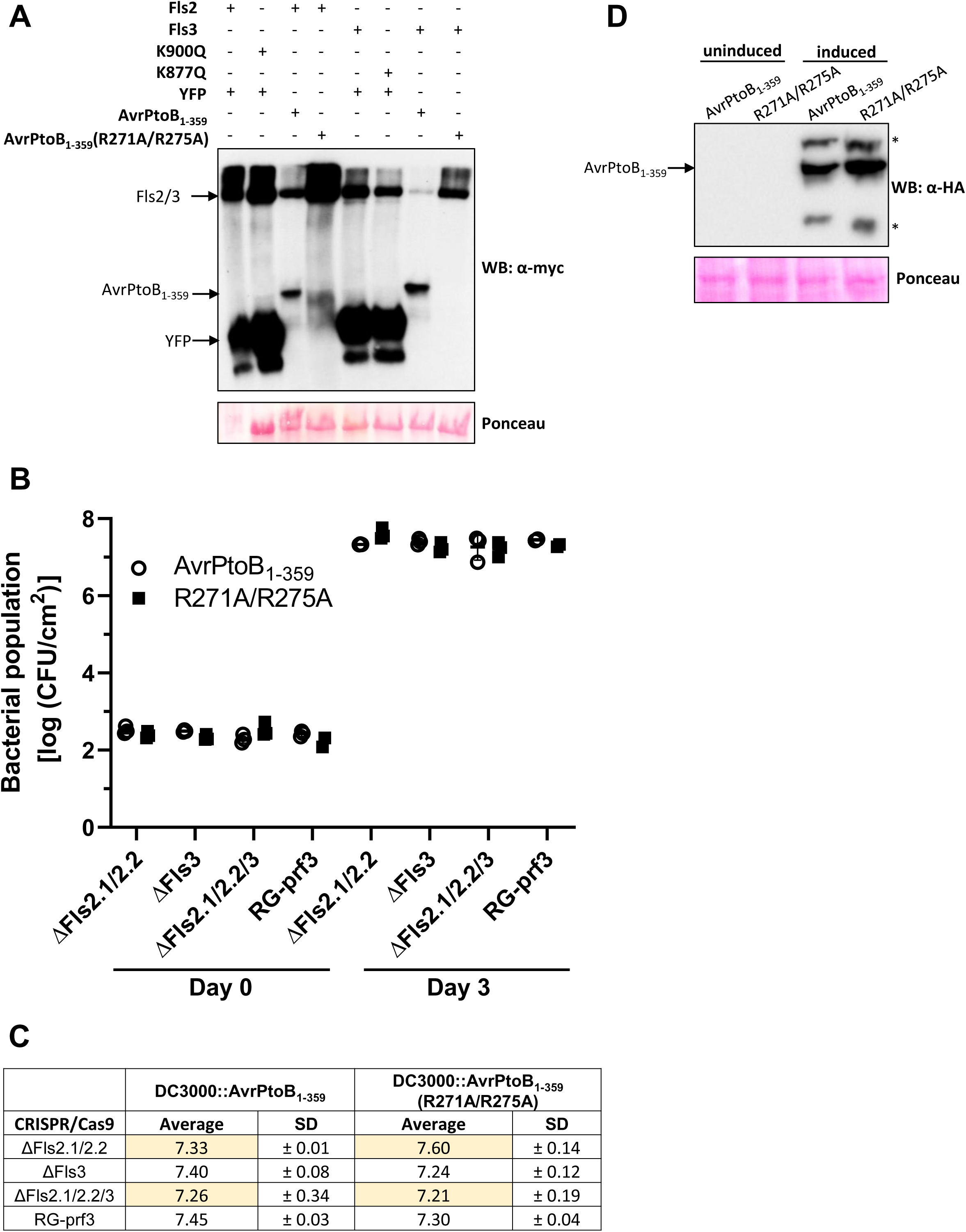
Supplementary information related to Figure 4. **(A)** Immunoblots showing protein expression of constructs in Figure 4A. 10μg of total protein extracted from *N. benthamiana* leaves co-agroinfiltrated and overexpressed with Fls2, Fls3, Fls2(K900Q), Fls3(K887Q), or YFP and AvrPtoB_1-359_ or AvrPtoB_1-359_(R271A/R275A) was run on SDS-PAGE and immunoblotted using an anti-cMyc antibody. Ponceau stain of the PVDF membrane was performed to compare loading. Constructs were overexpressed under a 35S promoter. **(B)** Bacterial populations in CRISPR/Cas9 mutant tomato plants *Fls2.1* and *Fls2.2* (ΔFls2.1/2.2), *Fls3* (ΔFls3), or *Fls2.1, Fls2.2*, and *Fls3* (ΔFls2.1/2.2/3) or wild type Rio Grande-prf3 (RG-prf3). Bacteria were measured 0 and 3 days after vacuum-infiltrating plants with bacterial suspensions (1 x 10^4^ CFU/mL) using a DC3000 strain expressing AvrPtoB_1-359_ (labeled AvrPtoB_1-359_) or its variant AvrPtoB_1-359_(R271A/R275A) (labeled R271A/R275A). Shown are the means of three individual plants indicated as separate points, and horizontal lines are the means of the three plants +/- s.d.. There were no statistically significant differences using a pairwise t-test in bacterial growth between the two strains for each plant (*P*>0.05). **(C)** Table showing the averages and standard deviations (SD) of the data in **(B)**. **(D)** Immunoblot showing expression of AvrPtoB_1-359_ and AvrPtoB_1-359_(R271A/R275A) expressed from a plasmid in DC3000 before and after induction of the hrp inducible promoter in minimal media. The C-terminal HA tag was detected using an anti-HA antibody, and Ponceau staining was performed to compare loading. *Protein of unexpected size specifically detected in AvrPtoB_1-359_-expressing bacteria, as previously reported in Cheng *et al* (2011).

**Figure S5.**
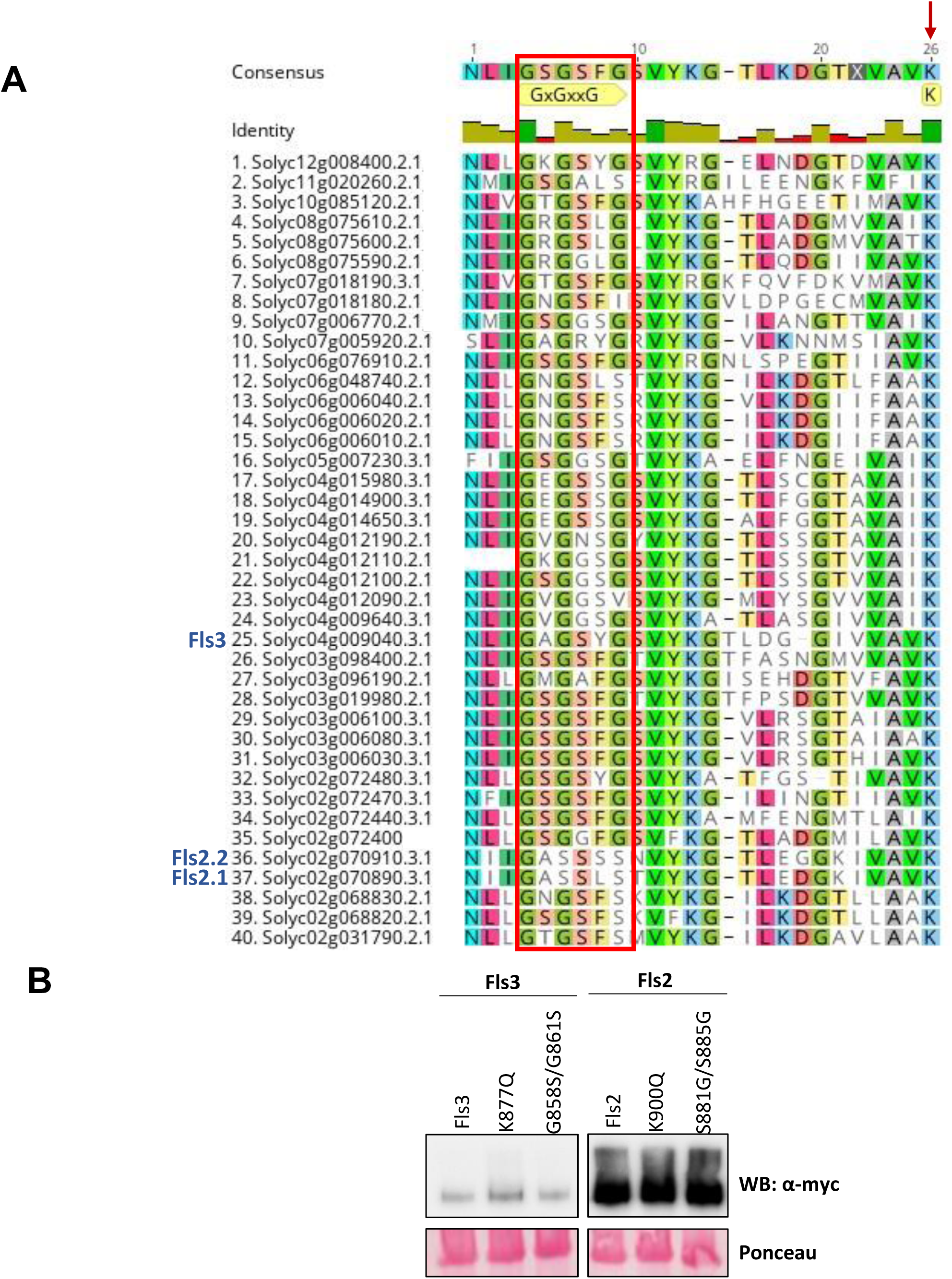
Supplementary information related to Figure 6. **(A)** Alignment of subdomain I of the RLKs present in class IX-a. The GxGxxG motif and the lysine (K) involved in ATP binding are annotated in yellow. The GxGxxG motif is also boxed in red and the lysine is indicated with a red arrow. Identity is shown with green indicating conservation, olive indicating high identity, and pink indicating low identity. Of the 42 RLKs in the class, two genes (Solyc10g085110 and Solyc03g118330) do not encode a subdomain I region or have the active lysine residue and are not included in the alignment. The alignment was assembled using the Geneious R11 software. **(B)** Immunoblot showing expression of proteins related to Figure 6. 5μg of total protein was extracted from *N. benthamiana* leaves agroinfiltrated with each of the Fls3 and Fls2 constructs or their variants and run on SDS-PAGE. Proteins were detected using an anti-cMyc antibody and Ponceau staining of the PVDF membrane was performed to compare loading. Constructs were overexpressed under the control of the CaMV 35S promoter. All samples shown are from the same experiment.

**Figure S6.**
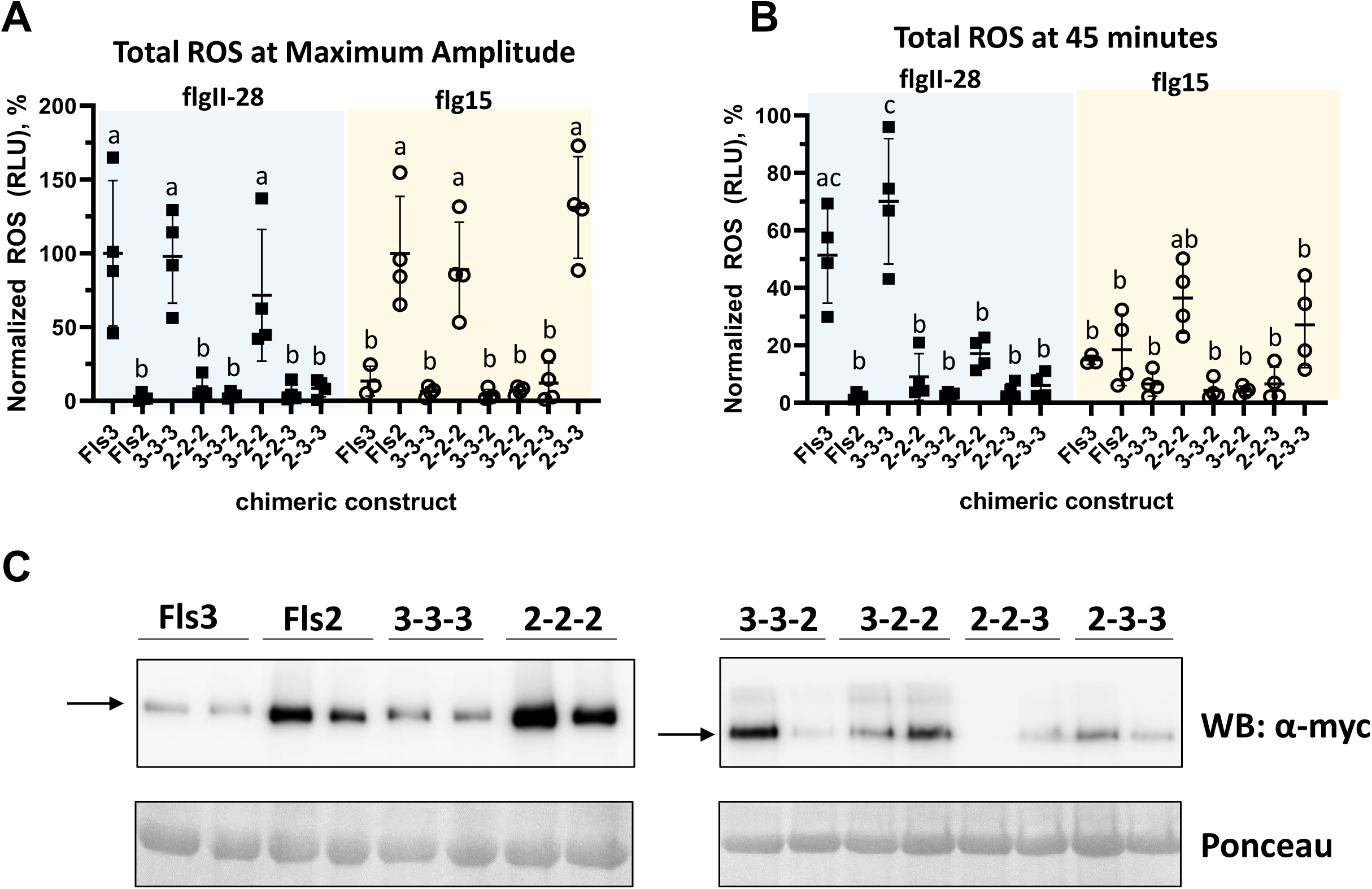
Only the chimeric constructs with a TM and KD originating from the same receptor respond to flgII-28 or flg15 peptide. Related to Figure 7. **(A)** Normalized total ROS response at the maximum amplitude for Fls3-flgII-28 or Fls2-flg15, 20 min (Fls3) or 12 min (Fls2) after peptide treatment (10nM flgII-28 or flg15). **(B)** Normalized total ROS response at 45 minutes after peptide treatment (10nM flgII-28 or flgII-15). Values in **(A)** and **(B)** were normalized to the maximum ROS (RLU) of the positive controls (Fls3 or Fls2) on each ROS plate. Constructs were transiently expressed in *N. benthamiana* under control of the CaMV 35S promoter. Data shown are the means of four individual plants indicated as separate points, and horizontal lines are the means of the four plants +/-s.d. (n=4 plants). Statistical significance was determined by Two-way ANOVA followed by Tukey’s multiple comparisons post-test. Data are representative of three independent experiments. Black squares signify samples treated with flgII-28, and open circles were samples treated with flg15. **(C)** Immunoblot showing expression of the chimeric constructs transiently expressed in *N. benthamiana*. Two samples from two individual plants (technical replicates) of the same experimental replicate (biological replicate) are shown for each infiltrated construct. 5μg of total protein was run on SDS-PAGE and proteins were detected using an anti-cMyc antibody.

**Table S1.**
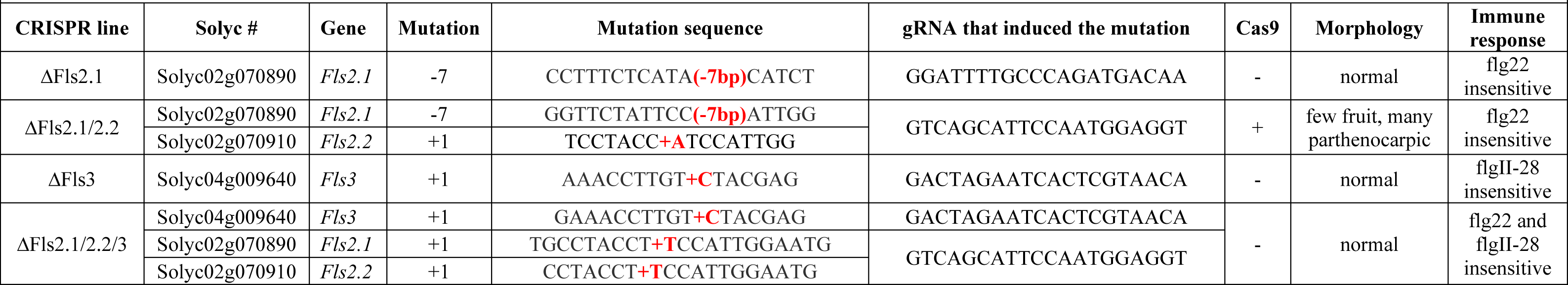
Generation of CRISPR/Cas9-mediated knockouts of the flagellin-sensing genes. Related to Figure 1 and Figure S1.

**Table S2.**
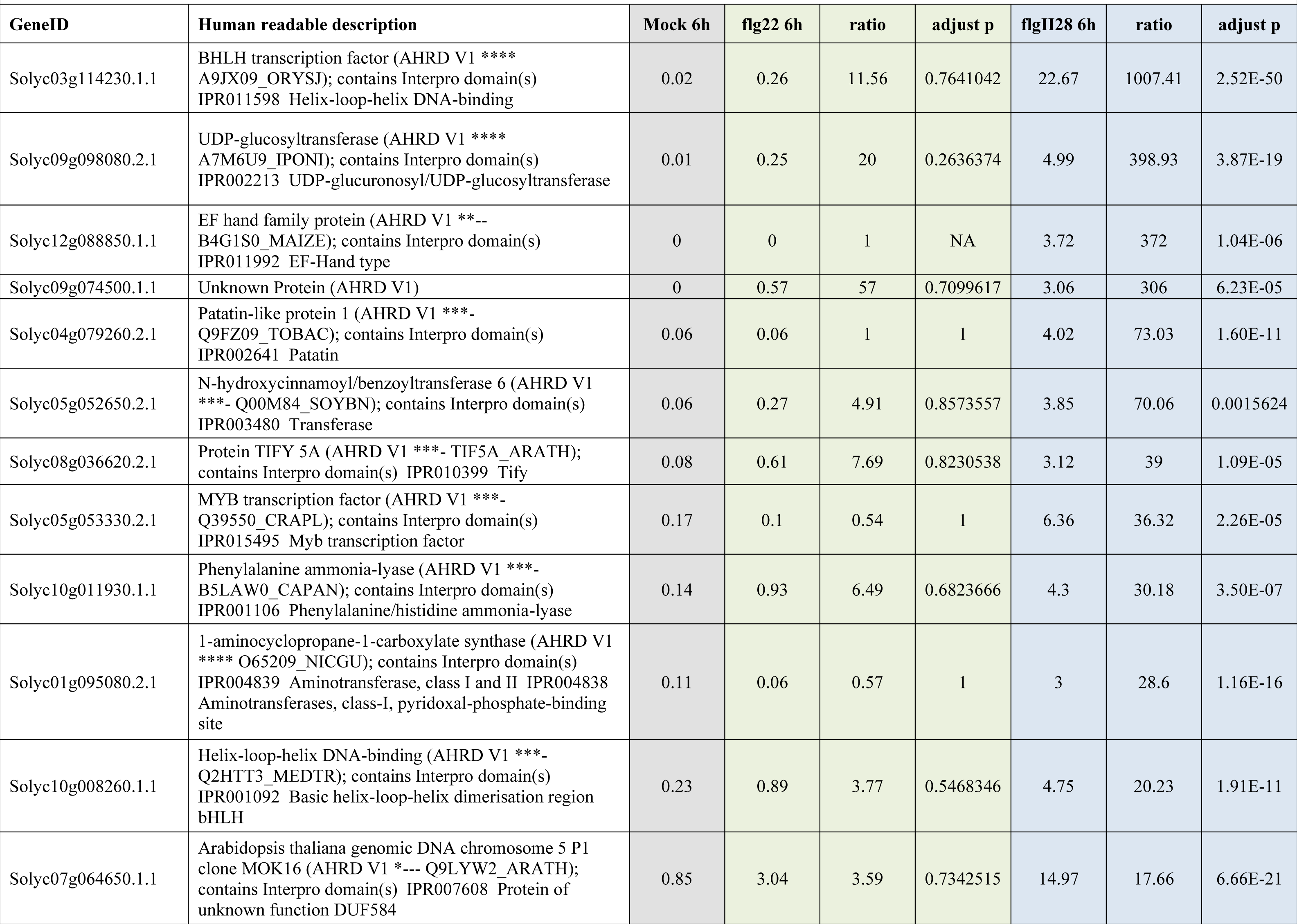
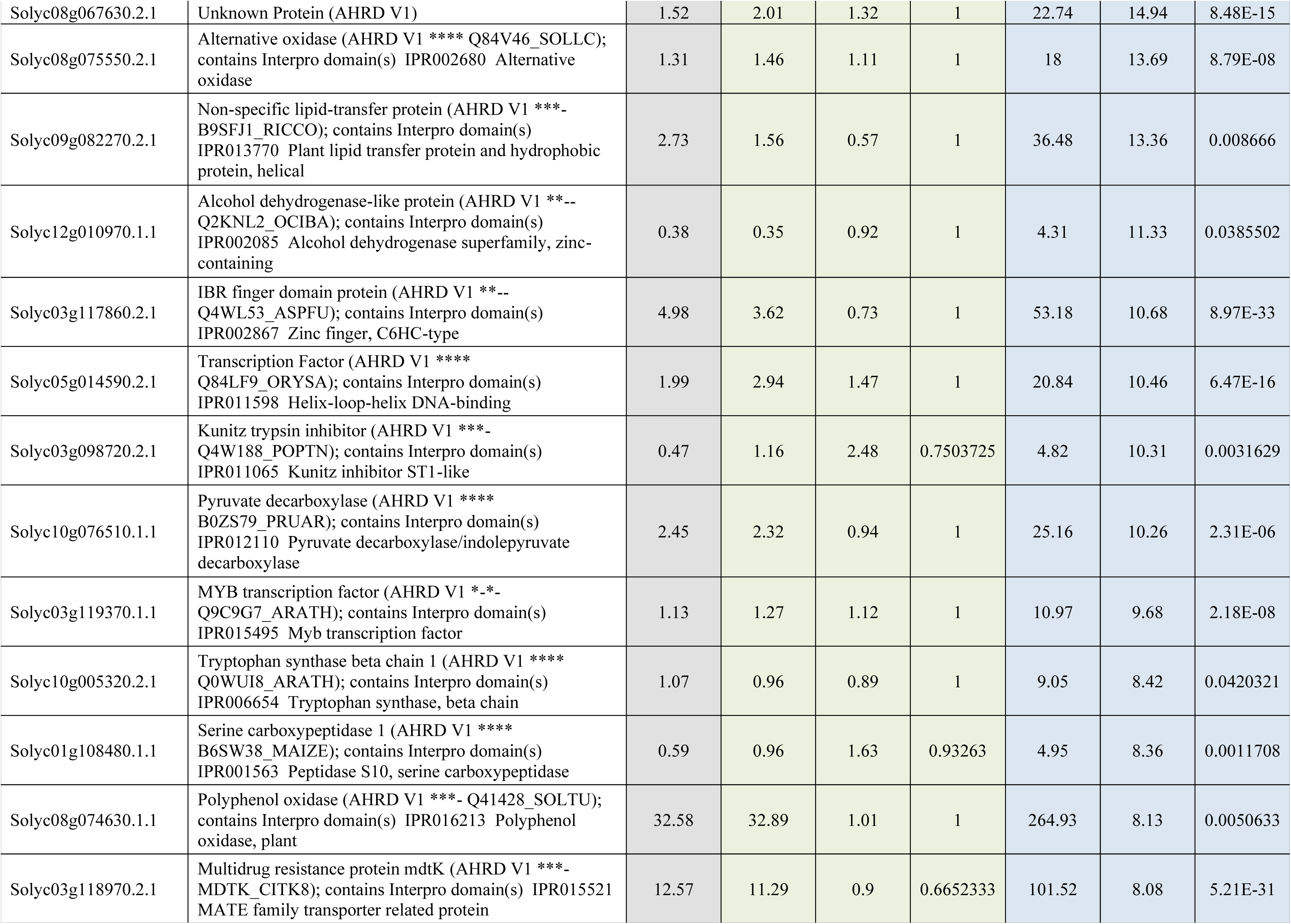
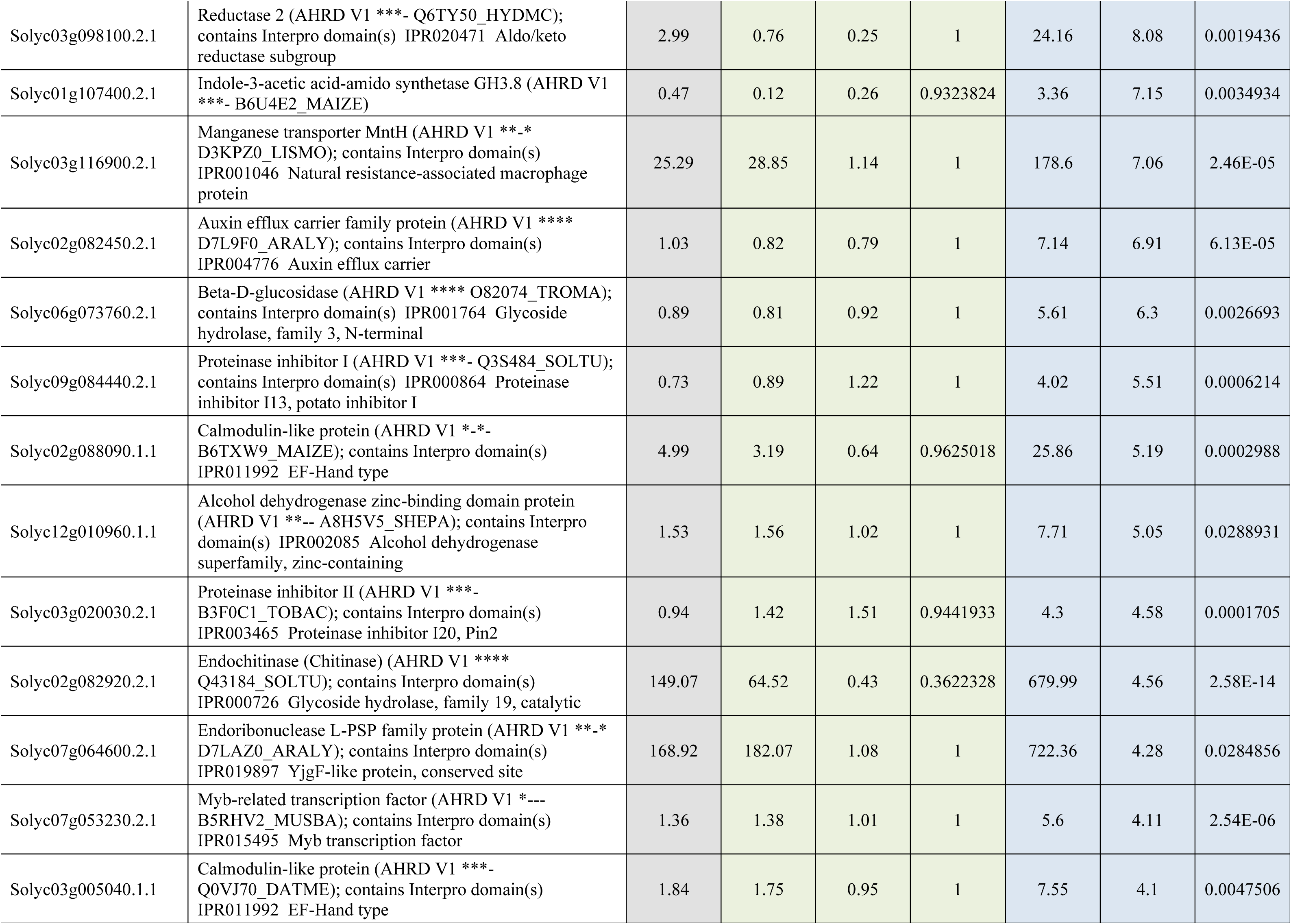
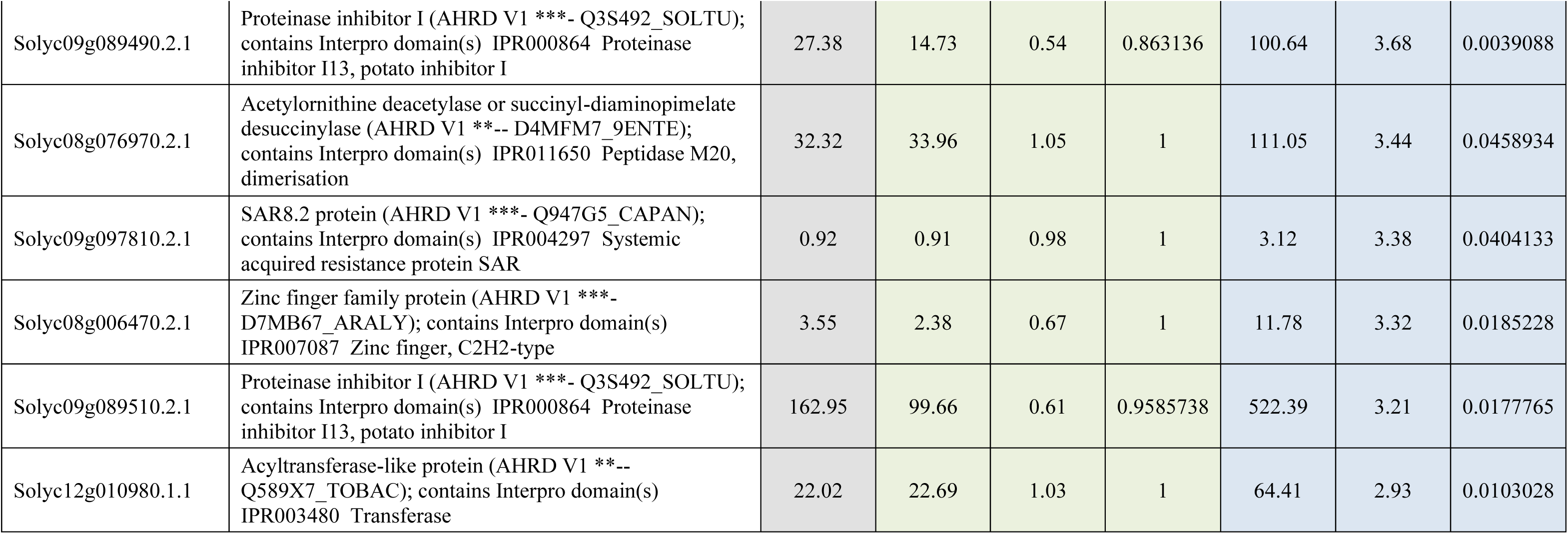
List of genes induced upon flgII-28 treatment by RNA-Seq (Pombo et al., 2017; Rosli et al., 2013). Related to Figure 3 and Table S3.

**Table S3.** GO Term analysis of flgII-28-induced genes from RNASeq (Pombo et al., 2017; Rosli et al., 2013) (all terms, q<0.05). Related to Figure 3 and Table S2.

|  |  |  |  |  |  |  |  |  |
| --- | --- | --- | --- | --- | --- | --- | --- | --- |
| GO.ID | Term | Annotated | Count | Expected | p-value | q-value | Aspect | Genes |
| GO:0030162 | regulation of proteolysis | 118 | 6 | 0.2 | 4.20E-08 | 7.79E-05 | P | Solyc03g020030.2, Solyc03g098720.2, Solyc03g117860.2, Solyc09g084440.2, Solyc09g089490.2, Solyc09g089510.2 |
| GO:0010466 | negative regulation of peptidase activity | 64 | 5 | 0.11 | 7.30E-08 | 7.79E-05 | P | Solyc03g020030.2, Solyc03g098720.2, Solyc09g084440.2, Solyc09g089490.2, Solyc09g089510.2 |
| GO:0010951 | negative regulation of endopeptidase activity | 64 | 5 | 0.11 | 7.30E-08 | 7.79E-05 | P | Solyc03g020030.2, Solyc03g098720.2, Solyc09g084440.2, Solyc09g089490.2, Solyc09g089510.2 |
| GO:0052547 | regulation of peptidase activity | 64 | 5 | 0.11 | 7.30E-08 | 7.79E-05 | P | Solyc03g020030.2, Solyc03g098720.2, Solyc09g084440.2, Solyc09g089490.2, Solyc09g089510.2 |
| GO:0052548 | regulation of endopeptidase activity | 64 | 5 | 0.11 | 7.30E-08 | 7.79E-05 | P | Solyc03g020030.2, Solyc03g098720.2, Solyc09g084440.2, Solyc09g089490.2, Solyc09g089510.2 |
| GO:0051346 | negative regulation of hydrolase activity | 69 | 5 | 0.12 | 1.10E-07 | 9.78E-05 | P | Solyc03g020030.2, Solyc03g098720.2, Solyc09g084440.2, Solyc09g089490.2, Solyc09g089510.2 |
| GO:0045861 | negative regulation of proteolysis | 76 | 5 | 0.13 | 1.80E-07 | 1.37E-04 | P | Solyc03g020030.2, Solyc03g098720.2, Solyc09g084440.2, Solyc09g089490.2, Solyc09g089510.2 |
| GO:0032269 | negative regulation of cellular protein metabolic process | 145 | 5 | 0.25 | 4.40E-06 | 2.67E-03 | P | Solyc03g020030.2, Solyc03g098720.2, Solyc09g084440.2, Solyc09g089490.2, Solyc09g089510.2 |
| GO:0051248 | negative regulation of protein metabolic process | 146 | 5 | 0.25 | 4.50E-06 | 2.67E-03 | P | Solyc03g020030.2, Solyc03g098720.2, Solyc09g084440.2, Solyc09g089490.2, Solyc09g089510.2 |
| GO:0051336 | regulation of hydrolase activity | 207 | 5 | 0.36 | 2.40E-05 | 1.28E-02 | P | Solyc03g020030.2, Solyc03g098720.2, Solyc09g084440.2, Solyc09g089490.2, Solyc09g089510.2 |
| GO:0032268 | regulation of cellular protein metabolic process | 370 | 6 | 0.64 | 3.30E-05 | 1.60E-02 | P | Solyc03g020030.2, Solyc03g098720.2, Solyc03g117860.2, Solyc09g084440.2, Solyc09g089490.2, Solyc09g089510.2 |
| GO:0051246 | regulation of protein metabolic process | 379 | 6 | 0.65 | 3.70E-05 | 1.64E-02 | P | Solyc03g020030.2, Solyc03g098720.2, Solyc03g117860.2, Solyc09g084440.2, Solyc09g089490.2, Solyc09g089510.2 |
| GO:0043086 | negative regulation of catalytic activity | 234 | 5 | 0.4 | 4.40E-05 | 1.81E-02 | P | Solyc03g020030.2, Solyc03g098720.2, Solyc09g084440.2, Solyc09g089490.2, Solyc09g089510.2 |
| GO:0031324 | negative regulation of cellular metabolic process | 400 | 6 | 0.69 | 5.00E-05 | 1.88E-02 | P | Solyc03g020030.2, Solyc03g098720.2, Solyc08g036620.2, Solyc09g084440.2, Solyc09g089490.2, Solyc09g089510.2 |
| GO:0044092 | negative regulation of molecular function | 243 | 5 | 0.42 | 5.30E-05 | 1.88E-02 | P | Solyc03g020030.2, Solyc03g098720.2, Solyc09g084440.2, Solyc09g089490.2, Solyc09g089510.2 |
| GO:0009611 | response to wounding | 148 | 4 | 0.25 | 0.00011 | 3.67E-02 | P | Solyc08g036620.2, Solyc09g084440.2, Solyc09g089490.2, Solyc09g089510.2 |
| GO:0004866 | endopeptidase inhibitor activity | 67 | 5 | 0.13 | 1.60E-07 | 1.10E-04 | F | Solyc03g020030.2, Solyc03g098720.2, Solyc09g084440.2, Solyc09g089490.2, Solyc09g089510.2 |
| GO:0061135 | endopeptidase regulator activity | 67 | 5 | 0.13 | 1.60E-07 | 1.10E-04 | F | Solyc03g020030.2, Solyc03g098720.2, Solyc09g084440.2, Solyc09g089490.2, Solyc09g089510.2 |
| GO:0030414 | peptidase inhibitor activity | 68 | 5 | 0.13 | 1.70E-07 | 1.10E-04 | F | Solyc03g020030.2, Solyc03g098720.2, Solyc09g084440.2, Solyc09g089490.2, Solyc09g089510.2 |
| GO:0061134 | peptidase regulator activity | 68 | 5 | 0.13 | 1.70E-07 | 1.10E-04 | F | Solyc03g020030.2, Solyc03g098720.2, Solyc09g084440.2, Solyc09g089490.2, Solyc09g089510.2 |
| GO:0004867 | serine-type endopeptidase inhibitor activity | 42 | 4 | 0.08 | 1.10E-06 | 5.70E-04 | F | Solyc03g020030.2, Solyc09g084440.2, Solyc09g089490.2, Solyc09g089510.2 |
| GO:0004857 | enzyme inhibitor activity | 224 | 5 | 0.42 | 6.00E-05 | 2.59E-02 | F | Solyc03g020030.2, Solyc03g098720.2, Solyc09g084440.2, Solyc09g089490.2, Solyc09g089510.2 |

**Table S4.**
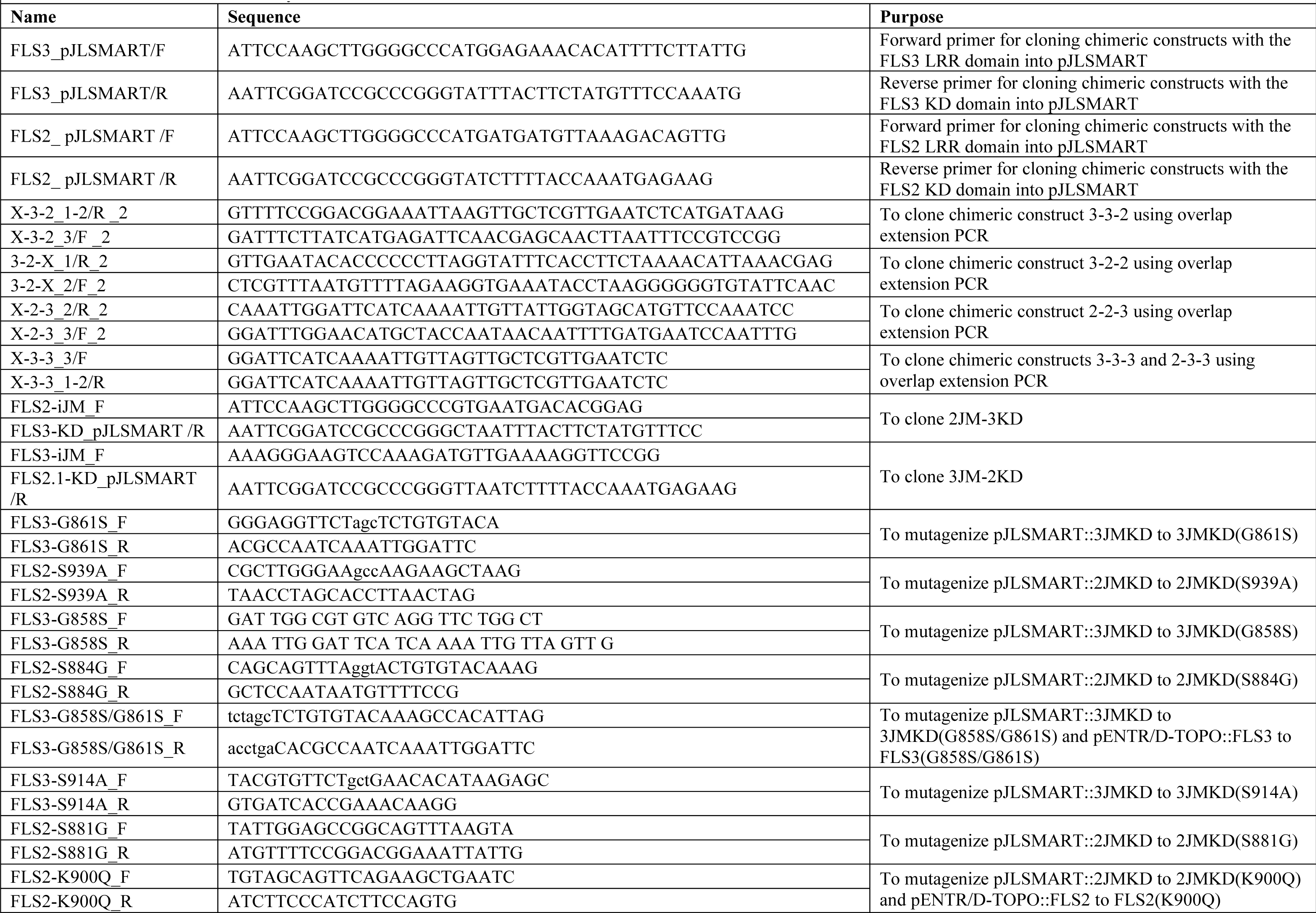
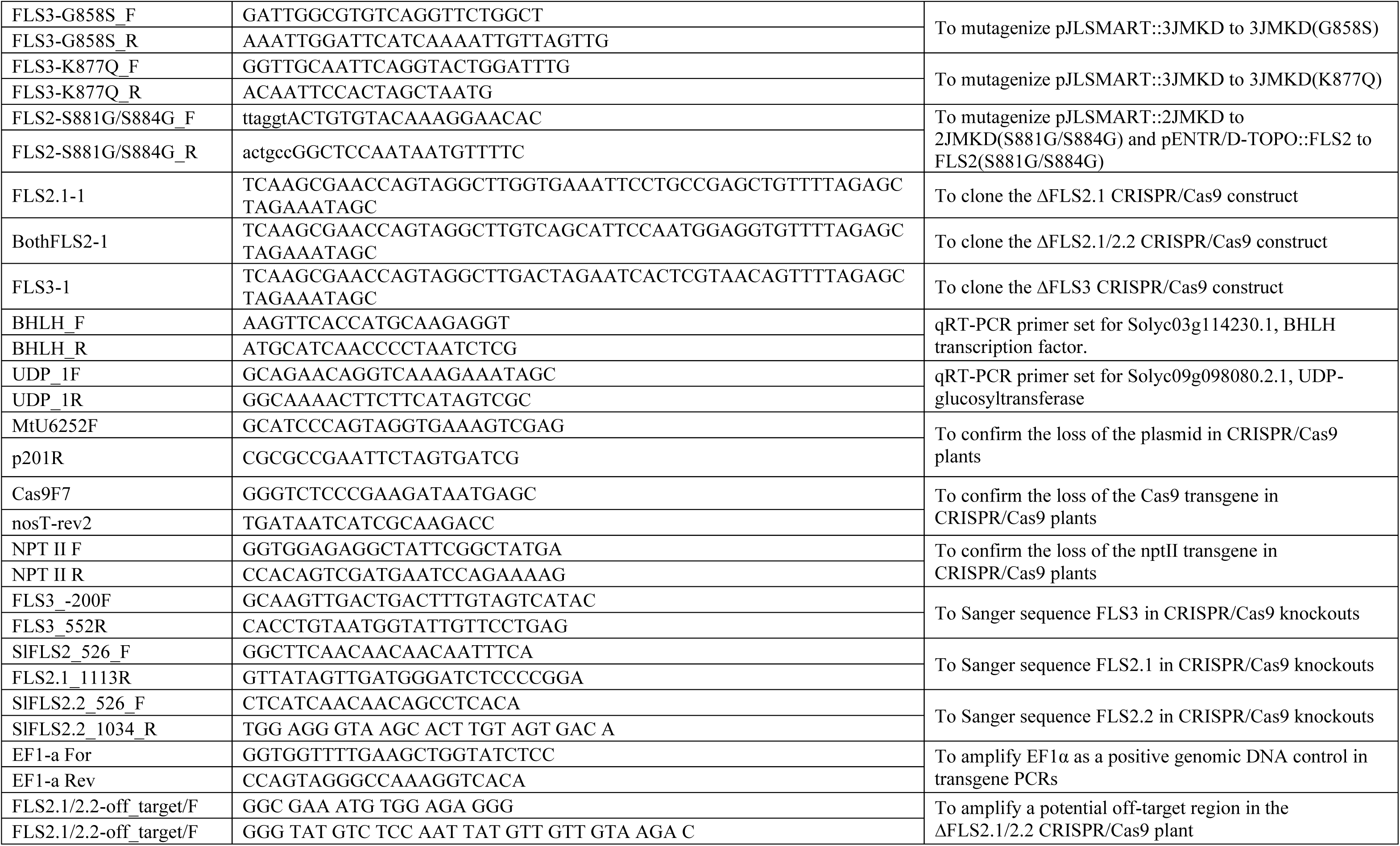
Primers used in this study. Related to the Methods section.

**Table S5.**
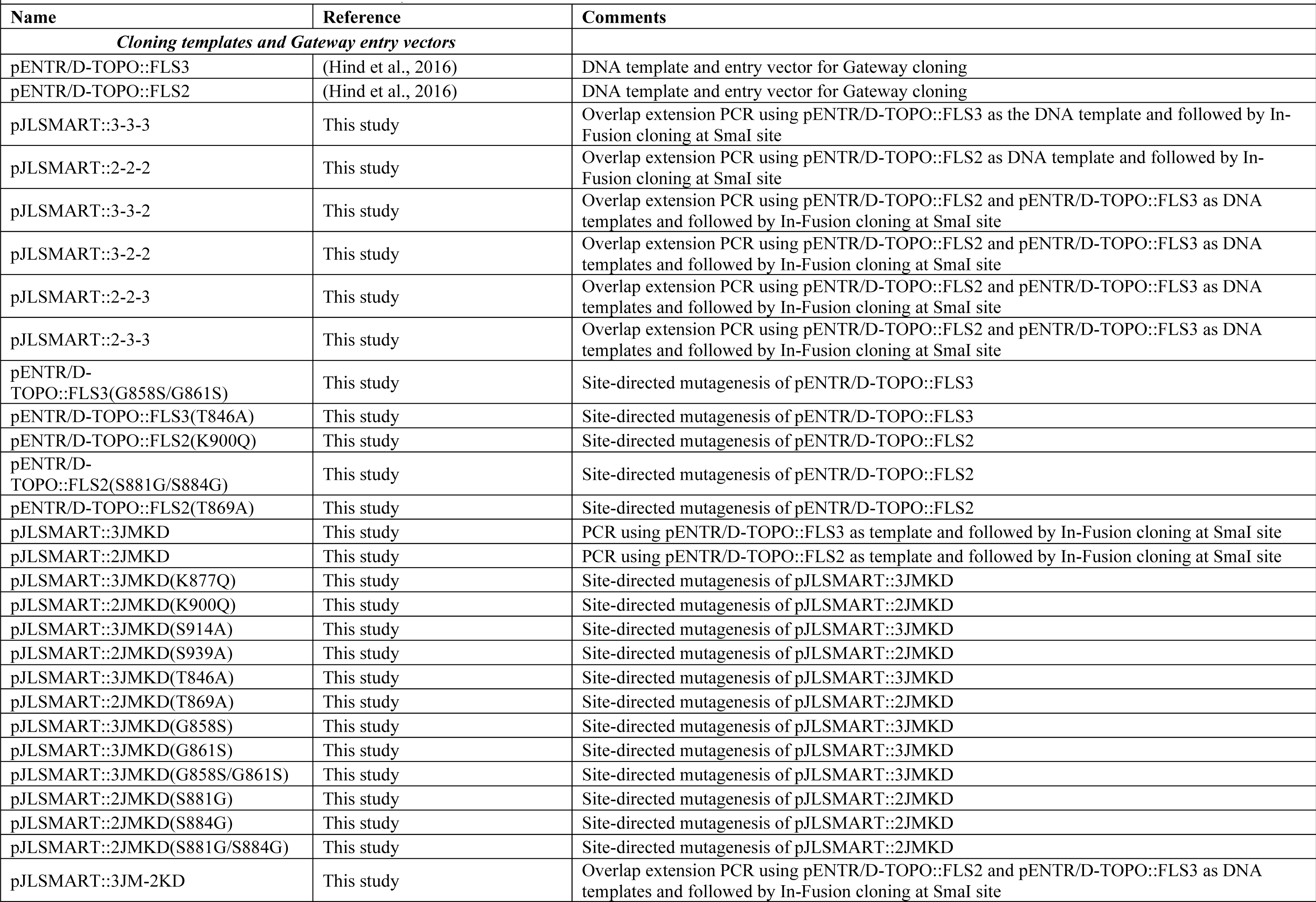
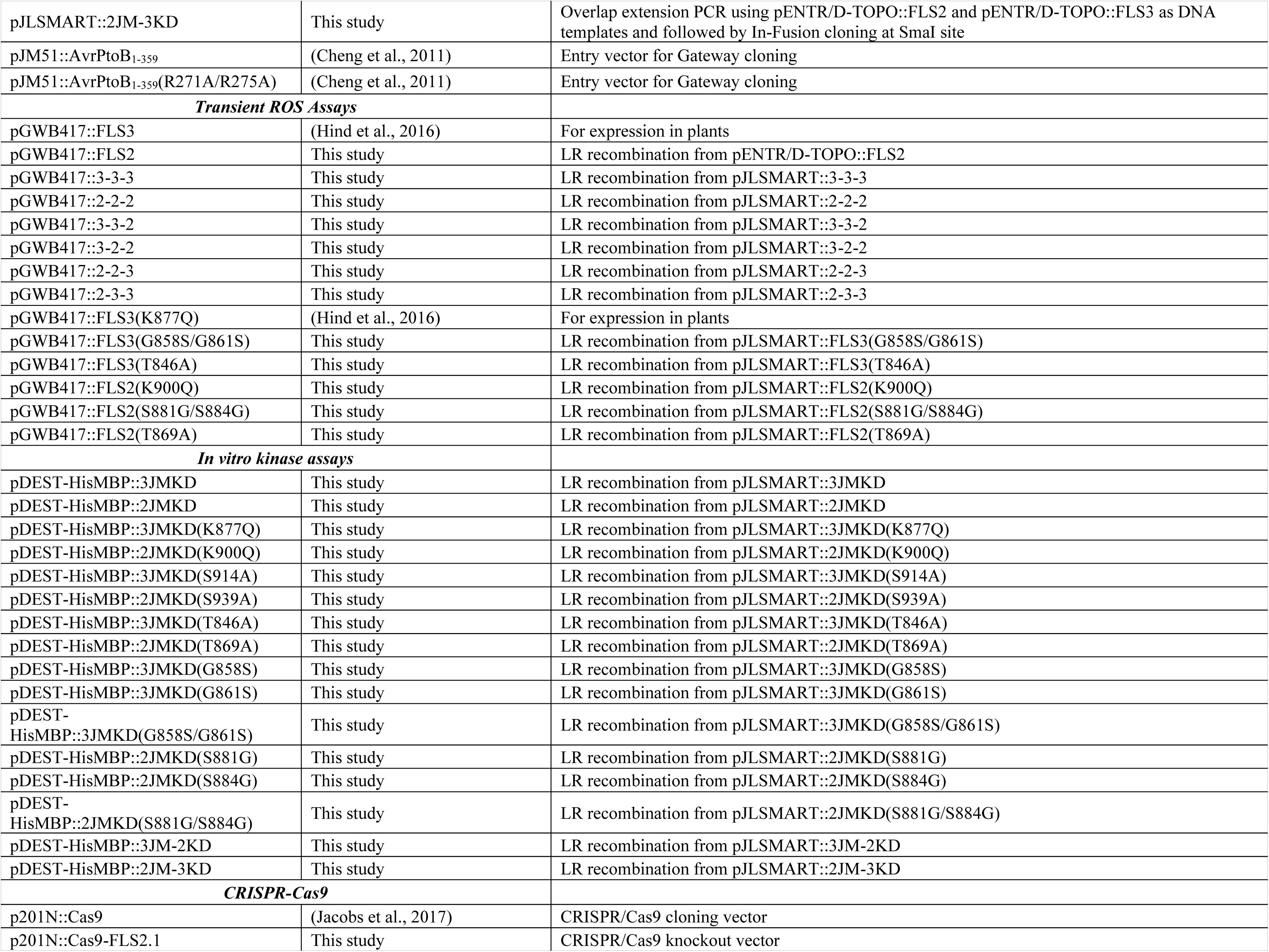
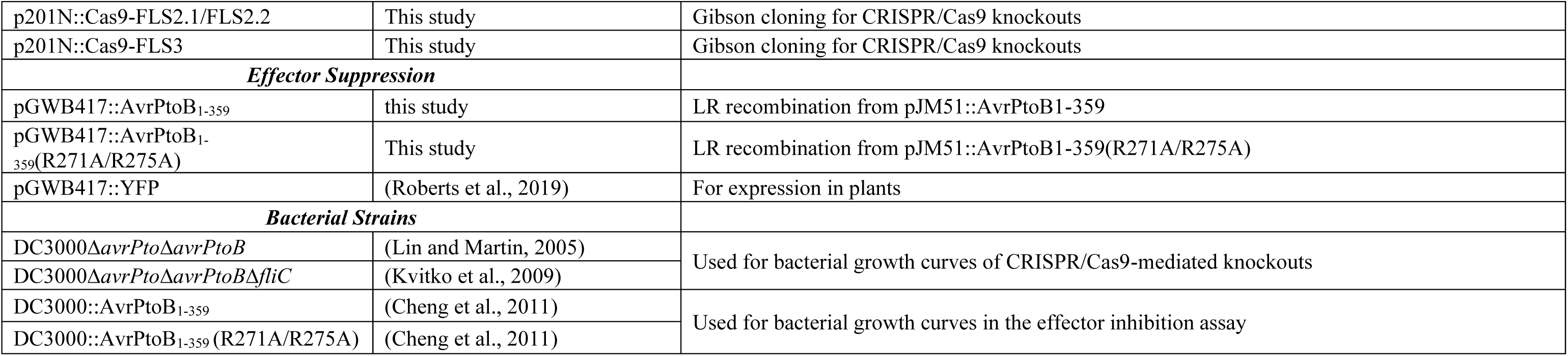
Constructs and strains used in this study. Related to the Methods section.

## Supplemental Methods

### Plant growth conditions, inoculations, and bacterial growth assays

Tomato seedlings were grown under the conditions described in (Roberts et al., 2019). *P. syringae* strains were grown on King’s B medium and suspended to a final concentration of 2 x 10^8^ CFU ml^-1^ for the dip inoculations or 1 x 10^4^ CFU ml^-1^ for the vacuum infiltrations as described previously (Roberts et al., 2019), and Silwet-L77 was added to a final concentration of 0.04%. Vacuum infiltrations were performed as previously described (Roberts et al., 2019). For the dip inoculations, three-week-old seedlings were placed in a 100% relative humidity chamber for 14 hours prior to inoculation, then dipped into the bacterial suspension for 10 seconds. Seedlings were placed back into the humidity chamber for 2 hours after inoculation. Leaflets were then excised from the plants and surface disinfected for two minutes in half-strength bleach before sampling, and bacterial populations were quantified on Day 0 and Day 2 or Day 3 as described in (Roberts et al., 2019). Experiments were repeated three times (n=3 plants) and shown is a single representative replicate. Data are means of individual plants in a single replicate (represented as points) and horizontal lines are the means of the three plants in a single experiment. Error bars are +/-s.d. and significance was determined using the Prism 8 program (GraphPad Software, https://www.graphpad.com/scientific-software/prism/). For a list of bacterial strains used in this study, see Table S3.

### Generation of CRISPR Cas9-mediated knockout lines

Geneious R9 software (https://www.geneious.com/) was used to design gRNAs to target *Fls3*, *Fls2.1*, or both *Fls2.1*/*Fls2.2* as described previously (Jacobs et al., 2017; Zhang et al., 2020) using the tomato genome version SL2.5 (Tomato Genome Consortium, 2012). The gRNAs selected for each of the genes were: *Fls3*, TGTTACGAGTGATTCTAGTC; *Fls2.1*, GAAATTCCTGCCGAGCTGGG; *Fls2.1/Fls2.2*, TACCTCCATTGGAATGCTGAC. Possible off-targets were predicted using the Geneious software. The gRNAs were cloned into the p201N::Cas9 vector using the Gibson assembly kit (New England Biolabs, www.neb.com), transformed into *Agrobacterium tumafaciens* strain LBA4404, and transformed into tomato (Rio Grande-prf3) as described in (Jacobs et al., 2017; Zhang et al., 2020). To induce mutations in both *Fls3* and *Fls2.1/2.2* in the same plant (ΔFls2.1/2.2/3), the constructs used to induce the individual mutations were transformed into LBA4404 and the Agrobacterium cultures were mixed 1:1 prior to tomato transformation. Tomato transformations were performed at the Boyce Thompson Institute transformation facility (Gupta and Van Eck, 2016; Van Eck et al., 2019). More information may be found in Table S1–3 and Fig. S1, and on the Plant CRISPR database at plantcrispr.org.

### Peptides used in the ROS bioassays

The flgII-28 (ESTNILQRMRELAVQSRNDSNSSTDRDA), flg22 (QRLSTGSRINSAKDDAAGLQIA), or *E. coli* flg15 (RINSAKDDAAGLQIA) peptides were commercially ordered from GenScript (https://www.genscript.com/) with >95% purity. Solubility testing was performed by the company, and all peptides were soluble in water. Peptides were also all predicted soluble using a prediction software (https://pepcalc.com/peptide-solubility-calculator.php). Multiple independent batches of flgII-28, flg22, and flg15 peptides were used throughout this paper, and the same observations were made with all batches. Because it is known that flg22 can be a contaminant in commercial peptides (Mueller et al., 2012), all peptides were confirmed not to be contaminated with flg22 before experimental use by testing them in ROS assays using Rio Grande, ΔFls2 plants (which do not respond to flg22 (Jacobs et al., 2017)), and Yellow Pear (which does not respond to flgII-28 (Hind et al., 2016)). New peptide batches were compared side-by-side with old batches in ROS assays to confirm the sensitivity was similar in Rio Grande. We did not observe any instances of contamination or changes in sensitivities between batches of peptides. The extended ROS response was always observed for flgII-28 in tomato and has been observed in previous studies of flgII-28 (Clarke et al., 2013; Hind et al., 2016). The extended ROS response to flgII-28 was observed regardless of peptide concentration (10 nM-1 μM).

### Cloning

The chimeric constructs were generated via overlap extension PCR using pENTR/D-TOPO::Fls3 and pENTR/D-TOPO::Fls2 as templates (Hind et al., 2016). The 2KD, 2JMKD, 3KD, and 3JMKD ORFs were amplified from pENTR/D-TOPO::Fls3 or pENTR/D-TOPO::Fls2 using PCR. PCR products were inserted into the entry vector pJLSMART (Mathieu et al., 2014) at the SmaI cut site using the In-Fusion cloning kit and following the manufacturer’s instructions (Takara, https://www.takarabio.com/). Sequences were confirmed via Sanger sequencing, and then the chimeric construct ORFs were recombined into the Gateway vector pGWB417 (Nakagawa et al., 2009; Nakagawa et al., 2007) using the LR Clonase II following the manufacturer’s instructions (Thermo Fisher Scientific, https://www.thermofisher.com/us/en/home.html). Recombination was confirmed using Sanger sequencing. Mutagenesis of the 2JMKD, 3JMKD, Fls2, and Fls3 clones was performed using the entry vectors and the Q5 Site-Directed Mutagenesis kit following the manufacturer’s instructions (New England Biolabs, www.neb.com). After mutations were confirmed using Sanger sequencing, ORFs were recombined into pDEST-HisMBP (2JMKD or 3JMKD) (Nallamsetty et al., 2005) or pGWB417 (Fls2 or Fls3) using the LR Clonase II enzyme. For a full list of primers and constructs used in this study, see Table S2 and S3.

### Agroinfiltrations

Binary vectors harboring Fls3 or Fls2 were transformed into *Agrobacterium tumefaciens* strain GV3101. Cultures were grown in induction medium for 5 hours at room temperature with shaking before adjusting the concentrations for infiltration. All cultures were prepared to a final OD_600_ of 0.2 and, the Fls3- and Fls2-containing bacterial cultures were mixed 1:1 with *Agrobacterium* cultures expressing the *p19* viral suppressor of silencing. For Figure 4, the bacterial cultures containing Fls3 or Fls2 and AvrPtoB_1-359_ or AvrPtoB_1-359_(R271A/R275A) were mixed in equal parts with the *p19* (1:1:1) before infiltration.

